# STAT5 Gain-of-Function Variants Promote Precursor T-Cell Receptor Activation to Drive T-Cell Acute Lymphoblastic Leukemia

**DOI:** 10.1101/2022.12.21.519945

**Authors:** Tobias Suske, Helena Sorger, Frank Ruge, Nicole Prutsch, Mark W. Zimmerman, Thomas Eder, Barbara Maurer, Christina Wagner, Susann Schönefeldt, Katrin Spirk, Alexander Pichler, Tea Pemovska, Carmen Schweicker, Daniel Pölöske, Dennis Jungherz, Tony Andreas Müller, Myint Myat Khine Aung, Ha Thi Thanh Pham, Kerstin Zimmel, Thomas Krausgruber, Christoph Bock, Mathias Müller, Maik Dahlhoff, Auke Boersma, Thomas Rülicke, Roman Fleck, Patrick Thomas Gunning, Tero Aittokallio, Satu Mustjoki, Takaomi Sanda, Sylvia Hartmann, Florian Grebien, Gregor Hoermann, Torsten Haferlach, Philipp Bernhard Staber, Heidi Anne Neubauer, Alfred Thomas Look, Marco Herling, Richard Moriggl

**Author notes:** Corresponding author: Richard Moriggl University of Veterinary Medicine Vienna Veterinärplatz 1 1210 Wien, Austria.

## Abstract

T-cell acute lymphoblastic leukemia (T-ALL) is an aggressive immature T-cell cancer. Hotspot mutations in JAK-STAT pathway members *IL7R*, *JAK1* and *JAK3* were analyzed in depth. However, the role of *STAT5A* or *STAT5B* mutations promoting their hyperactivation is poorly understood in the context of T-cell cancer initiation and acute leukemia progression. Importantly, the driver mutation *STAT5B^N642H^* encodes the most frequent activating STAT5 variant in T-ALL associated with poor prognosis. Here, we show that hyperactive STAT5 promotes early T-cell progenitor (ETP)-ALL-like cancer in mice and upregulated genes involved in T-cell receptor signaling (TCR), even in absence of surface TCR promoting. Importantly, these genes were also overexpressed in human T-ALL and other STAT5-dependent T-cell cancers. Moreover, human T-ALL cells were sensitive to pharmacologic inhibition by dual STAT3/5 degraders or ZAP70 tyrosine kinase blockers. Thus, we define STAT5 target genes in T-ALL that promote pre-TCR signaling mimicry. We propose therapeutic targeting using selective ZAP70 or STAT3/5 inhibitors in a subgroup of T-ALL patients with prominent IL-7R-JAK1/3-STAT5 activity.

**Significance:** We provide detailed functional characterizations of hyperactive STAT5A or STAT5B in thymic T-cell development and transformation. We found that hyperactive STAT5 transcribes T-cell-specific kinases or pre-TCR signaling hubs to promote T-ALL. Biomolecular and next-generation-sequencing methods, transgenesis and pharmacologic interference revealed that hyperactive STAT5 is a key oncogenic driver that can be targeted in T-ALL using STAT3/5 or SYK family member tyrosine kinase inhibitors.

**Conflict of interest:** The authors declare no potential conflicts of interest.

## Introduction

T-ALL represents up to 15% of pediatric and ∼25% of adult ALL cases (1,2). The disease is characterized by the malignant expansion of immature T-cells harboring chromosomal rearrangements leading to activation of oncogenes, including transcription factors like TAL1/2, TLX1/3, LMO1/2 or HOXA (3,4). These rearrangements often involve translocations to TCR gene enhancers, promoting their aberrant expression (3). This induces self-renewal, T-cell differentiation arrest and transformation through accumulation of further mutations (5,6), like gain-of-function (GOF) *NOTCH1* mutations, losses of tumor suppressor gene functions (e.g. *PTEN*, *CDKN2A, TP53*) or inactivation of genes encoding epigenetic regulators (e.g. *EZH2*, *SUZ12*, *EED*, 3,4,7). Furthermore, GOF mutations in *IL7R*, *JAK1*, *JAK3* or *STAT5B* are found in >30% of pediatric/adult T-ALL cases (4,8–11). Particularly, serine-to-cysteine substitutions in the extracellular juxtamembrane domain of the interleukin 6 receptor (IL-7R) α chain, causing disulfide bridge formation without ligand binding, or mutations in the transmembrane domain promoting homodimerization of the IL-7Rα chain, are well-characterized and known to induce constitutive JAK1-STAT5 activation (12–14). Moreover, overexpression of IL-7R or inactivation of phosphatases such as CD45 can lead to activation of the IL-7R-JAK1/3-STAT5B pathway in T-ALL (8,12). STAT5A and STAT5B are strongly related, showing ∼92% amino acid identity and ∼94% similarity (15). STAT5B is described as a stronger oncoprotein compared to STAT5A, but both display redundant transcriptional activity through heterodimerization (16–19). Importantly, *STAT5B^N642H^* is a recurrent, activating hotspot mutation found in T-cell cancers, including several mature leukemias and lymphomas, as well as NK-cell and myeloid neoplasms such as eosinophilia or acute myeloid leukemia (AML) (16,20–23). The aggressive nature of aberrant STAT5 activity is illustrated by transgenic mice expressing STAT5B^N642H^ or STAT5A^S710F^, which develop peripheral T-cell lymphoma/leukemia (PTCL), displaying dominant differentiation towards CD8^+^ effector memory T-cells (17–19). STAT5B^N642H^ is also found in T-ALL or in phenotypically similar T-cell lymphoblastic lymphoma (T-LBL), both diseases primarily affecting children (4,16,19,24,25). Here, we use T-ALL as terminology and include T-LBL as well, further supported by the fact that no clinical distinction between T-ALL and T-LBL exists for murine tumors. The presence of STAT5B^N642H^ was found to correlate with worse prognosis and a higher incidence of relapse upon chemotherapy in a cohort of 301 T-ALL patients (24).

Complete loss of STAT5 is embryonically lethal because of defective erythropoiesis in mice with pure C57BL/6 or BALB/c background. However, the few (∼2%) homozygous STAT5-null survivor mice on a mixed Sv129/C57BL/6 background displayed ∼98% reduction in thymocyte numbers (26). This emphasizes the fact that both STAT5A/B are essential for T-cell expansion and survival during mammalian development, in particular for T-cell differentiation (26). Several studies have revealed that the IL-7R-JAK1/3-STAT5 axis is essential for development of different thymic T-cell subsets, inducing transcription of cell cycle progression genes, BCL-2 family members or transcription factors such as RUNX3 or FOXP3 (27,28). Particularly, common γ-chain (γ_c_) signaling governs CD8^+^ lineage commitment via STAT5, and its conditional deletion in DP thymocytes reduced the frequency of SP8 cells (29). However, it is currently unknown, how *STAT5A* or *STAT5B* GOF mutations affect thymic development and, ultimately, induce transformation of immature T-cells to T-ALL in presence or absence of TCR surface chain expression.

Here, we show that mutation-driven hyperactivation of STAT5A or STAT5B promotes T-ALL without the requirement to express surface TCR. Especially STAT5B^N642H^ causes drastic changes in thymic maturation, architecture and global gene transcription. Mechanistically, we found that hyperactive STAT5 drives expression of kinases that are involved in TCR signaling promoting an activated T-cell phenotype. STAT5B^N642H^ or STAT5A^S710F^ act both as oncoproteins in a redundant fashion to induce T-ALL, but the clinically relevant STAT5B^N642H^ causes more aggressive phenotypes. We suggest that these findings are also highly relevant for other T-cell cancers. Blocking STAT3/5 and ZAP70 individually or in combination impaired T-ALL growth/survival. We conclude that STAT5 and T-cell receptor ligated tyrosine kinases represent valuable pharmacologic targets in T-ALL with high IL-7R-JAK1/3-STAT5 activation.

## Results

### STAT5B^N642H^ is recurrently mutated in T-ALL patients and alters T-cell development

*NOTCH1* GOF mutations are dominant in T-ALL patients, but interestingly *IL-7R-JAK1/3-STAT5B* axis GOF mutations are also found in up to 30% of patients (3). We data-mined eight independent studies with sequencing data from 1234 T-ALL patients (4,11,24,25,30–33) and found that a mean of 3.8% (0.6-12.9%) of patients harbor *STAT5B* mutations (**Fig. 1A**). *STAT5B^N642H^* was the most frequent recurrent mutation, accounting for 38 of the 62 (61.3%) cases with *STAT5B* mutations (**Fig. 1B**). 5 out of 14 different *STAT5B* mutations have been detected in ≥3 T-ALL patients (**Fig. 1B**) and these patients had almost twice as high white blood cell (WBC) counts as patients with unmutated *STAT5B*, suggesting that STAT5B hyperactivation accelerates T-ALL cell growth and progression (**Fig. 1C**).

**Figure 1.**
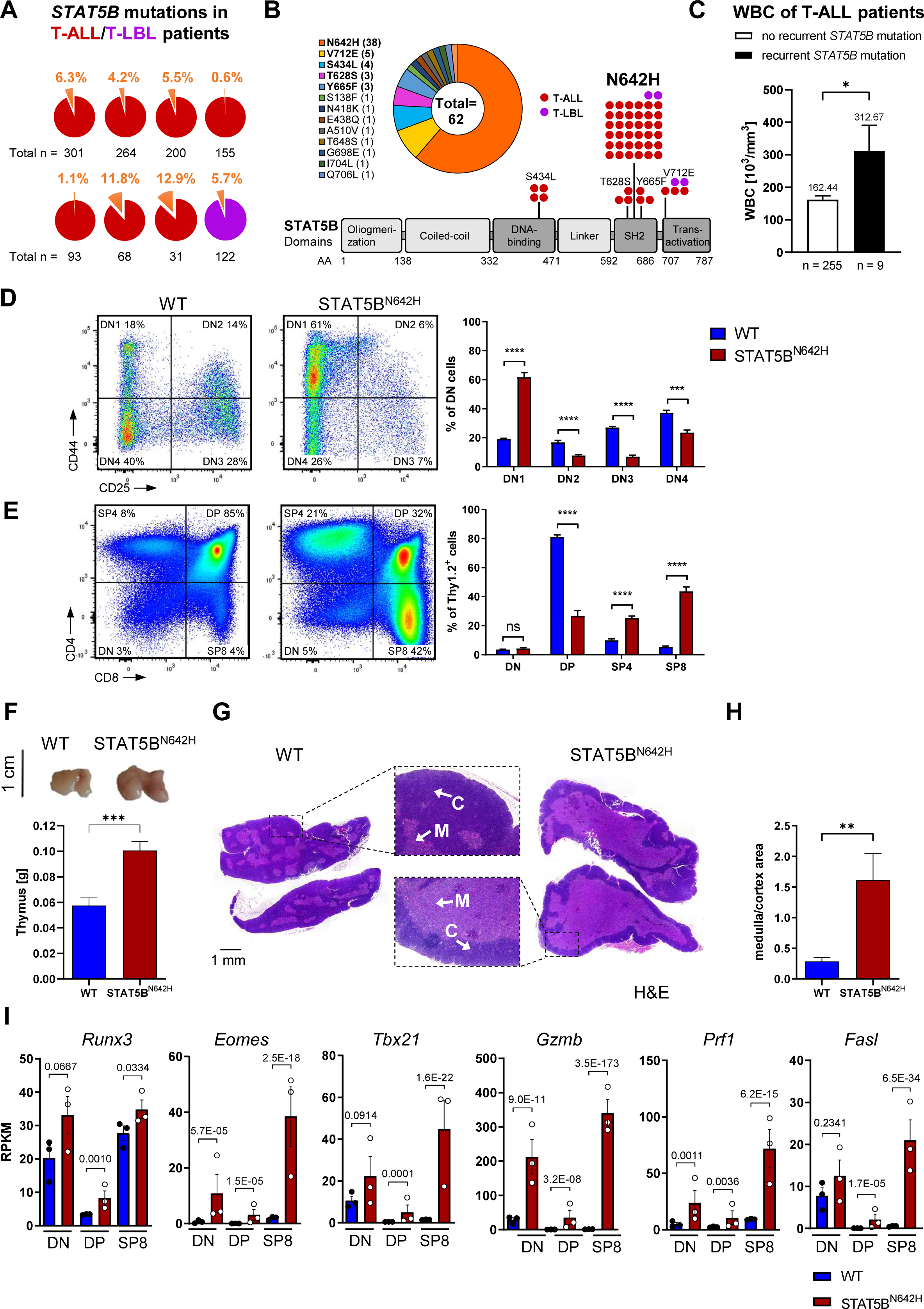
STAT5B^N642H^ occurs in T-ALL patients and impacts thymic development and architecture. **A**, Percentages of patients with somatic mutations in *STAT5B* found in 8 studies in T-ALL or T-LBL, total number of patients indicated below pie charts. **B**, Incidence and localization in the STAT5 protein with amino acid (AA) position of respective somatic mutations in *STAT5B* found in 1234 T-ALL or T-LBL patients from the studies indicated in **A**. **C**, White blood cell (WBC) numbers of patients with or without three or more *STAT5B* mutations indicated in **B**. Data were extracted from reference (4). **D** and **E**, Flow cytometry analysis and percentages of DN1, DN2, DN3, DN4 (CD25/CD44) and DN, DP, SP4, SP8 (CD4/CD8) cells from STAT5B^N642H^ mice (n = 7) and WT littermates (n = 7) gated on Thy1.2^+^ thymocytes. **H**, Representative images and thymus weights in g of eight-week-old WT (n = 7) or STAT5B^N642H^ (n = 7) mice. **I**, Representative hematoxylin and eosin (H&E) stained whole thymi from WT or STAT5B^N642H^ mice indicating cortical (C) and medullary (M) regions, original magnification (centered images): 10x, scale bar = 1 mm, and **J**, quantification of medullary-cortical area ratios thereof, with n (WT) = 6 and n (STAT5B^N642H^) = 7. **K**, RPKM values of *Runx3*, *Eomes*, *Tbx21*, *Gzmb*, *Prf1*, *Fasl* in DN, DP and SP8 stages of WT or STAT5B^N642H^ thymocytes. Experiments in **D**-**H** were performed twice independently. H&E-stainings in **I** of formalin-fixed and paraffin-embedded thymi from 7 mice of each genotype were assessed with ImageJ for cortical and medullary areas. In **A**, **D**-**H** and **J**, significant differences are indicated as \**P* < 0.05, \*\**P* < 0.01, \*\*\**P* < 0.001, \*\*\*\**P* < 0.0001, by unpaired two-tailed Student’s t-test. Error bars show mean +/-SEM. Error bars show mea +/-SEM. In **K**, adjusted *P* value (*Padj*) of respective comparisons were taken from DESeq2 analysis.

To *investigate* mechanistic consequences of STAT5B^N642H^ expression on T-cell transformation, we analyzed transgenic mice expressing human STAT5B^N642H^ under the control of the *Vav1*-promoter, driving transgene expression from the hematopoietic stem cell level onwards, including lymphoid progenitors (34). STAT5B^N642H^ mice succumb to disease at ∼10 weeks of age due to mature, post-thymic CD8^+^ T-cell neoplasia (17).

Thymic T-cell development undergoes CD4/CD8 double-negative (DN) to double-positive (DP) transition prior to CD4 or CD8 single-positive (SP4/SP8) lineage commitment (35). STAT5B^N642H^ mice exhibited progressing disease associated with increased spleen weight (**Supplementary Fig. S1A**). Increased STAT5 phosphorylation (pY-STAT5) in thymi of diseased STAT5B^N642H^ mice was confirmed by Western blotting (**Supplementary Fig. S1B**). Next, we determined DN1 (CD25^-^ CD44^+^), DN2 (CD25^+^ CD44^+^), DN3 (CD25^+^ CD44^-^), DN4 (CD25^-^ CD44^-^), DP, SP4 and SP8 populations by flow cytometry. We found that STAT5B^N642H^ promoted ∼3-fold expansion of the most immature DN1 cells, at the expense of DN2/3/4 populations compared to WT littermates at eight weeks of age (**Fig. 1D**). DP cells were 3-fold reduced, while SP4 and SP8 cells were ∼2.5- and ∼8-fold expanded in STAT5B^N642H^ mice (**Fig. 1E**). We found elevated levels of CD69 in DP thymocytes of STAT5B^N642H^ mice, suggesting an advantage in positive selection. Conversely, CD69 levels were reduced in SP4 and SP8 cells from STAT5B^N642H^, implying facilitated commitment to SP4 and SP8 lineages and a mature SP stage to facilitate thymic egress (36,37) (**Supplementary Fig. S1C**). Surface CD5 was reduced in STAT5B^N642H^ SP cells compared to WT controls (**Supplementary Fig. 1D**), which suggests a lower inhibitory effect on TCR signaling due to reduced binding to CD3ζ-ZAP70 components of the TCR/CD3 complex (38–40). These effects on thymic development in the STAT5B^N642H^ mice were already apparent at four and six weeks of age further accelerated upon disease progression (**Supplementary Fig. S1E**). Moreover, the thymic mass was elevated in STAT5B^N642H^ mice compared to WT littermates (**Fig. 1H**; **Supplementary Fig. S1F**). Maturation of T-cells is accompanied by migration from the thymic cortex to the medulla (35). H&E staining confirmed an increase in the medulla/cortex ratio in STAT5B^N642H^ thymi, in line with the expansion of SP T-cells (**Fig. 1I** and **J**). Next, we isolated CD8^+^ T-cells from STAT5B^N642H^ mice and transplanted them into Ly5.1^+^ recipients. Two- and eight-weeks post-transplant, Ly5.2^+^ CD8^+^ T-cells were present in high numbers in spleen, lymph nodes (LNs), bone marrow, blood, liver or lung, but lacking in thymus, confirming that changes in thymocyte ratios were thymus-intrinsic and not caused by infiltration of mature CD8^+^ T-cells (**Supplementary Fig. S1G**).

Surprisingly, the *Stat5a^S710F^* GOF mutation affected T-cell development milder compared to STAT5B^N642H^ mice at 28 weeks of age (**Supplementary Fig. S1H**), possibly due to the longer disease latency (25-45 weeks, 18). These differences highlight the impact of hyperactive STAT5 on T-cell development with expanded immature DN1 thymocytes and strongly increased SP8 thymocyte populations possibly culminating in T-ALL phenotypes as a result of increased positive selection associated with TCR signaling.

### Hyperactive STAT5A or STAT5B drive activated T-cell phenotypes in developing thymocytes

To understand thymic transcriptional reprogramming by hyperactive STAT5A/B, we collected DN, DP and the more dominantly expanded SP8 cells of diseased STAT5B^N642H^ and STAT5A^S710F^ and WT thymi by FACS-sorting and performed RNA-seq (Supplementary Fig. S2A and S2B). In WT DP thymocytes, *Stat5b* expression was much higher than other Stat mRNA, including *Stat5a*. (**Supplementary Fig. S2C** and **S2D**). The majority of differentially expressed genes in STAT5B^N642H^ versus WT thymocytes were upregulated, emphasizing the strong transcriptional activity of hyperactive STAT5B^N642H^ (**Supplementary Fig. S3A** and **S3B**). Mutually upregulated genes by STAT5B^N642H^ across all three T-cell subsets were enriched for GO-terms representing T-cell differentiation and activation, as well as chemotaxis, interferon type I/II signaling or host defense (**Supplementary Fig. S3C**). We found that mRNAs encoding for key transcription factors controlling CD8^+^ T-cell maturation, such as RUNX3, EOMES, T-bet and cytotoxic effector molecules Granzyme B, Perforin and Fas ligand – were highly upregulated throughout all T-cell developmental stages in STAT5B^N642H^ thymi (**Fig. 1K**). Moreover, differentially expressed genes (STAT5B^N642H^ vs. WT in DN, DP and SP8 subsets) correlated more with signatures of activated than resting T-cells (**Supplementary Table 1**). Differential expression patterns were conserved in sorted DN, DP and SP8 thymocytes from STAT5A^S710F^ mice (**Supplementary Fig. S4A-S4C**; **Supplementary Table 2**). Consistent with strong SP8 lineage commitment caused by hyperactive STAT5, purified STAT5B^N642H^ DP cells gave rise to SP8 lymphomas after transplantation into WT mice (**Supplementary Fig. S5A-S5B**). These data suggest that hyperactive STAT5 causes a pre-activation status that is independent of the T-cell development stage and TCR-mediated antigen signaling.

Normal WT T-cells display an activated state mainly in post-thymic, mature stages (41). Our RNA-seq profiles suggested that *STAT5A/B* GOF mutations induce the machinery for high-level activation in immature, unselected T-cells, possibly inducing transformation processes through upregulating T-cell activation signatures. Importantly, all analyzed thymic subsets, STAT5B^N642H^ and STAT5A^S710F^ induced transcriptional signatures correlating with human ETP-ALL with a reportedly high JAK/STAT activation (42) (**Supplementary Fig. S5D**).

Together, these results confirm that STAT5B^N642H^ enhances thymocyte CD8^+^ lineage commitment/expansion and that STAT5 GOF variants actively contribute to an activated T-cell phenotype and a T-ALL-like transcriptome.

### Hyperactive STAT5 oncoproteins induce thymic T-cell tumors upon arrest of T-cell development

Blocking immature stages of T-cell development and acquisition of self-renewal capacity through driver oncogenes is a hallmark of T-ALL (7). To prevent mature T-cell differentiation and to assess the ability of STAT5 GOF to induce immature, thymic T-cell leukemia, we crossed our hyperactive STAT5 mice and STAT5B WT controls with RAG2-deficient mice, in which T-cell development is blocked at the DN3 stage due to a lack of VDJ recombination and surface TCR αβ/γδ chain expressions(43) (**Fig. 2A**). Notably, in this setting we did not observe myeloid transformation and only one mouse of 30 developed a CD19^+^ B-cell lymphoma that was excluded from subsequent analysis (**Supplementary Fig. S6A**). All other STAT5 GOF RAG2^-/-^ mice developed a dominant T-ALL-like phenotype, irrespective of sex. STAT5B^N642H^ RAG2^-/-^ mice developed terminal disease between 15 and 22 weeks of age, which is a significantly longer latency than in STAT5B^N642H^ mice (**Fig. 2B**). In comparison, STAT5A^S710F^ RAG2^-/-^ mice succumbed to disease significantly later (25-45 weeks) than STAT5B^N642H^ RAG2^-/-^ mice, similar to STAT5A^S710F^ mice. The disease in STAT5 GOF RAG2^-/-^ mice was characterized by a massive increase in thymus/spleen weights and WBC counts compared to RAG2^-/-^ littermates, where thymus enlargement was most drastic, also compared to WT and STAT5B^N642H^ mice (**Fig. 2C** and **D**; **Supplementary Fig. S6B-S6D**). An increase in the numbers of Thy1.2-expressing cells by flow cytometry and high CD3 expression assessed by immunohistochemistry indicated strong thymic T-cell expansion in STAT5 GOF RAG2^-/-^ mice, despite the loss of VDJ recombination (**Supplementary Fig. S6E** and **S6F**), coupled with T-cell infiltration into the lung (**Fig. 2E**). In contrast, human wild type (h) STAT5B RAG2^-/-^ mice did not show any sign of disease when sacrificed at 35 weeks of age, similar to RAG2^-/-^ and WT control mice (**Fig. 2B**; **Supplementary Fig. S6G**).

**Figure 2.**
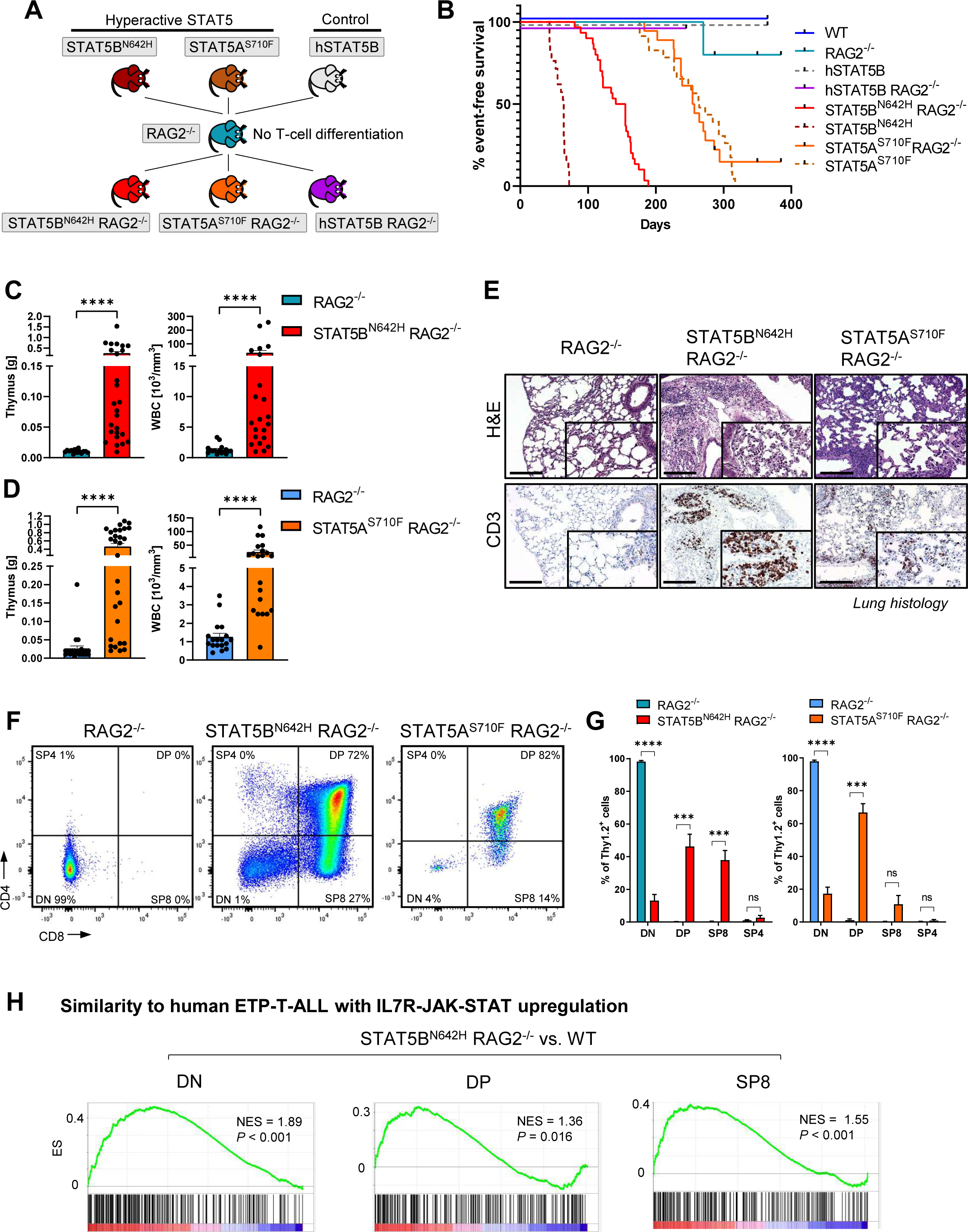
CD8+ lineage commitment and activated state of T-cells induced by STAT5BN642H. STAT5B^N642H^ induces immature thymic T-cell neoplasia in a RAG2^-/-^ background. **D, E,** GSEA comparing top 250 upregulated genes in human ETP-ALL (10) to deregulated genes in STAT5BN642H or STAT5AS710F DN, DP or SP8 vs. respective WT populations, NES: normalized enrichment score. **A**, Crossing scheme of different STAT5B or STAT5A mutant or WT mice with RAG2^-/-^ mice. **B**, Kaplan-Meier event-free survival plot of all mouse strains of indicated genotypes. **C**, Thymus weights in g of and WBC in 10^3^/mm^3^ of STAT5B^N642H^ RAG2^-/-^ (n = 26 and 22) compared to RAG2^-/-^ littermates (n = 15 and 16) and **D**, STAT5A^S710F^ RAG2^-/-^ (n = 29 and 21) compared to RAG2^-/-^ littermates (n = 25 and 18). **E**, H&E and immunohistochemical anti-CD3 stainings of lung tissues of STAT5 GOF RAG2^-/-^ and RAG2^-/-^ mice, original magnification: 10x and 20x (insets), scale bar = 200 µm, representative of 4 biological replicates. **F** and **G**, Flow cytometry analysis and relative abundance of DN, DP, SP8 and SP4 cells by CD4/CD8 staining gated on Thy1.2 in STAT5B^N642H^ RAG2^-/-^ (n = 12) and RAG2^-/-^ (n = 6) littermates and STAT5A^S710F^ RAG2^-/-^ (n = 4) and RAG2^-/-^ (n = 4) littermates. In **C**, **D** and **G**, significant differences are indicated as \*\*\**P* < 0.001, \*\*\*\**P* < 0.0001, by a Mann-Whitney U test. Error bars show mean +/-SEM.

Next, we aimed to evaluate T-cell immunophenotypic surface marker expression in the thymus at terminal disease by flow cytometry. In line with previous studies, staining for CD4 and CD8 was absent in RAG2^-/-^ thymi. Strikingly, in all STAT5B^N642H^ RAG2^-/-^ and STAT5A^S710F^ RAG2^-/-^ mice, the majority of thymocytes was constituted by DP and SP8 cells, indicating that STAT5 hyperactivation overcomes the developmental block in the DN stage of RAG2^-/-^ thymocytes and induces expression of CD4 and CD8 (**Fig. 2F** and **G**). Thymocytes of hSTAT5B RAG2^-/-^ mice revealed no significant upregulation of CD4 or CD8 compared to RAG2^-/-^ littermates (**Supplementary Fig. S6H**). We also detected elevated thymus weights and expression of CD4 and CD8 in younger (10–14-week-old), non-terminally-diseased STAT5B^N642H^ RAG2^-/-^ mice (**Supplementary Fig. S6I** and **S6J**).

Next, we performed RNA-seq on FACS-sorted thymic DN cells of RAG2^-/-^ and DN, DP and SP8 cells of diseased STAT5B^N642H^ RAG2^-/-^ and STAT5A^S710F^ RAG2^-/-^ mice (**Supplementary Fig. S7A**). Global changes in gene expression appeared similar in STAT5B^N642H^ RAG2^-/-^ and STAT5A^S710F^ RAG2^-/-^ compared to RAG2^-/-^ thymocytes, again indicating overlapping functions of both hyperactive *STAT5A/B* gene variants (**Supplementary Fig. S7B** and **S7C**) in their capacity to transform immature T-cells. Importantly, the STAT5B^N642H^-driven thymic neoplasms transcriptionally correlated with human ETP-ALL (**Fig. 2H**), whereas this correlation was weaker for STAT5A^S710F^-driven thymic disease, except for DN cells (**Supplementary Fig. S8A**). Altogether, these results confirm that STAT5A^S710F^ and the clinically more relevant STAT5B^N642H^ are drivers for T-ALL questioning their targeting in T-ALL.

### Immature thymocytes of STAT5B^N642H^ RAG2^-/-^ mice transform to T-ALL

The increase in thymus size of diseased STAT5B^N642H^ RAG2^-/-^ mice was the most drastic difference compared to RAG2^-/-^, WT or STAT5B^N642H^ mice (**Supplementary Fig. S5D**), suggesting the development of immature T-cell neoplasia. While cortical and medullary regions were histologically distinguishable in WT and STAT5B^N642H^ thymi, only cortical regions were identified in RAG2^-/-^ or STAT5B^N642H^ RAG2^-/-^ thymi (**Supplementary Fig. S8B**). We found widespread, strong expression of the proliferation marker Ki67 (**Supplementary Fig. S8C**) and the clinically relevant T-ALL marker Terminal deoxynucleotide Transferase (TdT) in STAT5B^N642H^ RAG2^-/-^ thymi (**Supplementary Fig. S8D**). High levels of TdT indicated immature thymocyte expansion and a phenotype similar to ETP-ALL. This supports the above findings that STAT5B^N642H^ RAG2^-/-^ thymocytes remain immature.

Comparing RNA-seq data from four mouse genotypes (WT, STAT5B^N642H^, RAG2^-/-^, STAT5B^N642H^ RAG2^-/-^), we found elevated levels of *Pim1* exclusively in STAT5B^N642H^ RAG2^-/-^ thymocytes (**Supplementary Fig. S9A**), confirmed by Western blot (**Supplementary Fig. S9B**). PIM1 kinase is a well-described oncogenic, direct STAT5 target gene, being targeted in T-ALL (44,45). Our findings suggest that STAT5 GOF mutations promote immature T-ALL initiation and progression.

### STAT5B^N642H^ upregulates TCR pathway genes

Next, we addressed the molecular basis of immature T-cell neoplasm development. The absence of surface CD3 (sCD3) attenuates the transition from DN to DP T-cells, whereas CD3 stimulation or transgenic TCR expression induces a transition to the DP stage in RAG-deficient mice (46). DN cells were sCD3-negative as expected, but surprisingly, DP and SP8 cells from STAT5B^N642H^ RAG2^-/-^ thymi expressed elevated levels of sCD3, as a possible prerequisite for this TCR-driven progress in thymic development (**Fig. 3A**). Notably, the CD3ζ chain was expressed higher in STAT5B^N642H^ RAG2^-/-^ than in RAG2^-/-^ thymi (**Fig. 3B**).

**Figure 3.**
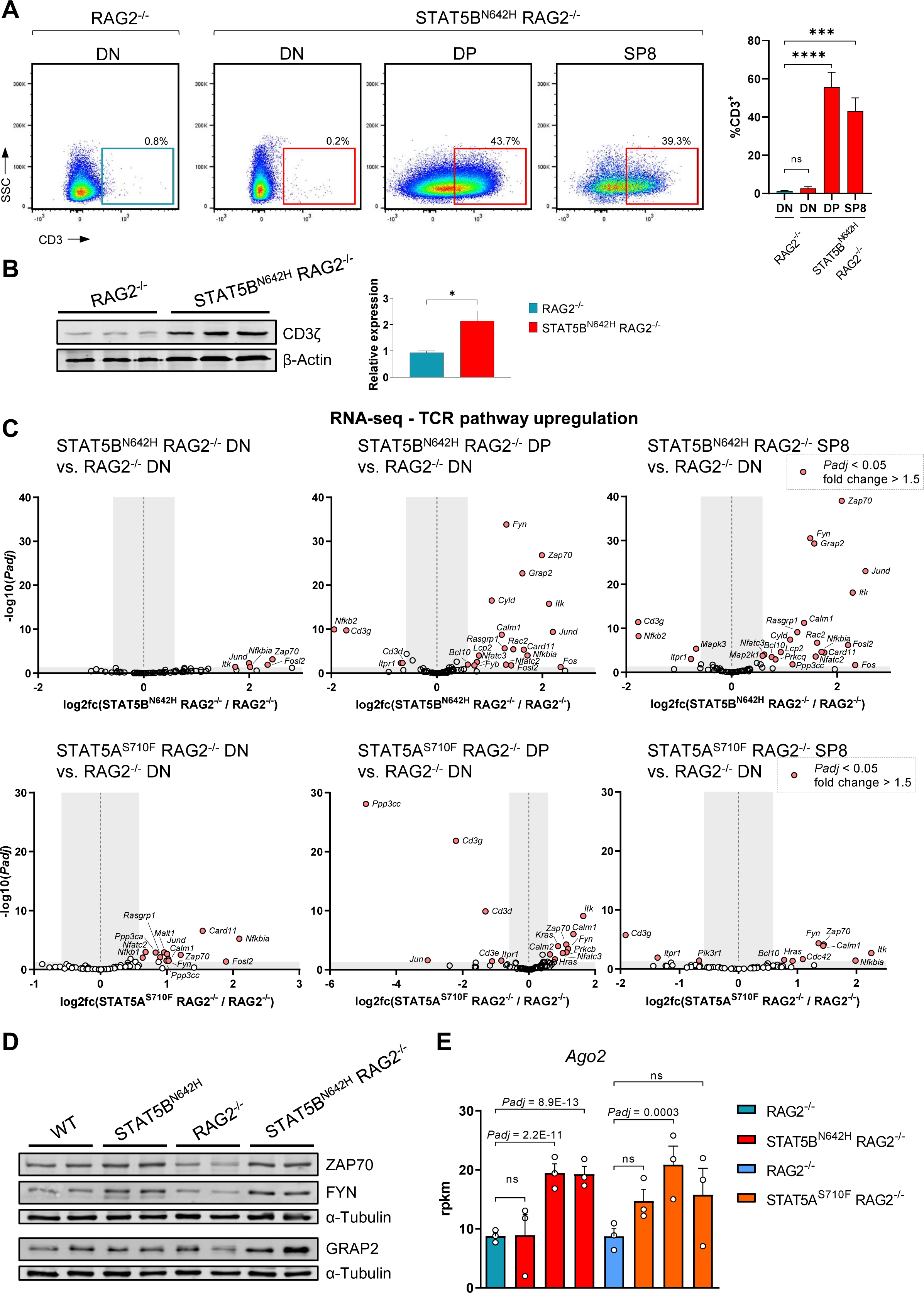
STAT5B^N642H^ RAG2^-/-^ T-ALL cells upregulate CD3 surface expression and TCR pathway genes. **A**, Flow cytometry analysis for CD3 surface expression in DN, DP or SP8 cells of thymi from STAT5B^N642H^ RAG2^-/-^ (n = 9) or RAG2^-/-^ (n = 5) littermates. **B**, Western blot analysis for CD3ζ expression in protein extracts of whole thymi of STAT5B^N642H^ RAG2^-/-^ or RAG2^-/-^ mice and quantification thereof, with n (STAT5B^N642H^ RAG2^-/-^) = 6 and n (RAG2^-/-^) = 4. **C**, Volcano-plots of RNA-seq data of DN, DP, SP8 (STAT5B^N642H^ RAG2^-/-^ and STAT5A^S710F^ RAG2^-/-^) vs. DN (respective RAG2^-/-^ littermates) cells, *Padj* and logfc determined by DESeq analysis. **D**, Western blot analysis for ZAP70, FYN and GRAP2 expression in protein extracts of whole thymi of indicated mouse genotypes. **E**, RPKM values of *Ago2* DN, DP, SP8 (STAT5B^N642H^ RAG2^-/-^ and STAT5A^S710F^ RAG2^-/-^) cells vs. DN cells (respective RAG2^-/-^ littermates), *Padj* determined by DESeq analysis. In **A** and **B**, significant differences are indicated as \**P* < 0.05, \*\*\**P* < 0.001, \*\*\*\**P* < 0.0001 by one-way Anova with Dunnet’s multiple comparison test (**A**) or unpaired two-tailed Student’s t-test (**B**). Error bars show mean +/- SEM. \**P* < 0.05, \*\**P* < 0.01 by unpaired two-tailed Student’s t-test. In **A** and **F**, error bars show mean +/-SEM.

Previous studies suggest that transition to the DP stage despite recombination deficiency can result from signaling downstream of the TCR pathway (46). Based on our above finding, we therefore more closely evaluated the transcription of genes involved in the TCR signaling cascade. Strikingly, STAT5B^N642H^ induced significant upregulation of 18 (DP) and 20 (SP8) genes of 71 genes encoding proteins that are implicated in TCR signaling (**Fig. 3C**). Fewer of these genes, were significantly deregulated in STAT5B^N642H^ RAG2^-/-^ DN cells, in line with weaker sCD3 expression in this subset (**Fig. 3C**). GSEA confirmed these results, where gene expression profiles of DP and SP8 cells correlated more with TCR pathway genes than DN cells (**Supplementary Fig. S9C**). Surprisingly, this signature was less pronounced in thymi of STAT5A^S710F^ RAG2^-/-^ mice, where only DN cells displayed moderate upregulation of the same genes (**Fig. 3C**). We confirmed that STAT5B^N642H^ RAG2^-/-^ thymocytes expressed higher levels of ZAP70 and FYN tyrosine kinases and the scaffold kinase adaptor GRAP2 protein (**Fig. 3D**; Supplementary Fig. S9D). Moreover, *Ago2* transcription was upregulated in the thymic subsets that also displayed upregulation of TCR pathway genes (**Fig. 3E**). While AGO2 is a regulator of miRNA processing, it has been recently attributed to non-canonical functions in TCR signaling upregulation by complex formation with e.g. ZAP70 (47). We conclude, that hyperactive STAT5 upregulates gene expression to mimic pre-TCR signaling encoding e.g. CD3ζ, AGO2, GRAP2 or kinases like ZAP70, FYN and ITK.

### Human T-ALL express high levels of TCR pathway genes

Next, we explored gene expression profiles of T-ALL patients to evaluate expression levels of TCR pathway genes. In three independent studies, *ZAP70*, *FYN*, *GRAP2, ITK* and *CD247* (encoding CD3ζ) were highly upregulated in T-ALL patients compared to healthy controls or other hematopoietic cancers, similar to our murine study (**Fig. 4A**; **Supplementary Fig. S10A**). Additionally, *LCK* as another key regulator of T-cell activation was highly upregulated in these datasets (**Fig. 4A**), although we did not observe significant *Lck* upregulation in our four neoplastic mouse models. Furthermore, we found that expression levels of *ZAP70*, *FYN*, *GRAP2*, *ITK*, *LCK* and two further TCR pathway kinases, *PLCG1* and *LAT* were the highest in human T-ALL cell lines compared to other hematopoietic cancer lines. (**Fig. 4B**; **Supplementary Fig. S10B**). Oncogenic TCR signaling has already been reported in PTCL (48–51) and higher expression levels of TCR pathway kinases in T-ALL compared to PTCL strengthen the hypothesis that they are key players in immature T-cell transformation. However, continuously high expression of these genes in PTCL confirm that TCR signaling might be targetable in a broad range of T-cell cancers. Examining publicly available CRISPR screening data, we found that in T-ALL cell lines, the dependency scores of *LCK*, *ZAP70* and *CD247* were higher than for other TCR pathway genes, reflecting that the CD3ζ-ZAP70 interaction could be a vulnerable node in T-ALL (**Supplementary Fig. S10C**). Moreover, high ZAP70 expression correlated with a higher risk of T-ALL relapse in patients (**Supplementary Fig. S10D**). We analyzed whole-exome-sequencing in conjunction with RNA-seq data of human T-ALL subtypes and found that immature T-ALL cells with *JAK/STAT* GOF mutations had increased *ZAP70* expression levels compared to unmutated ones (**Supplementary Fig. S10E**). Upon immunostaining of individual biopsies from T-ALL patients we detected ZAP70 activation by phosphorylation of tyrosine 319 (pY319) in more than 5% of stained tissue area in nine of ten samples, which further supports the concept of an activated T-cell signature in a subfraction of T-ALL cells in patient biopsies (**Fig. 4C** and **D**). Thus, our data indicate that active STAT5 or CD3ζ-ZAP70 signaling could be relevant targets in human T-ALL or possibly also several PTCL types, where STAT5B^N642H^ is also found (16).

**Figure 4.**
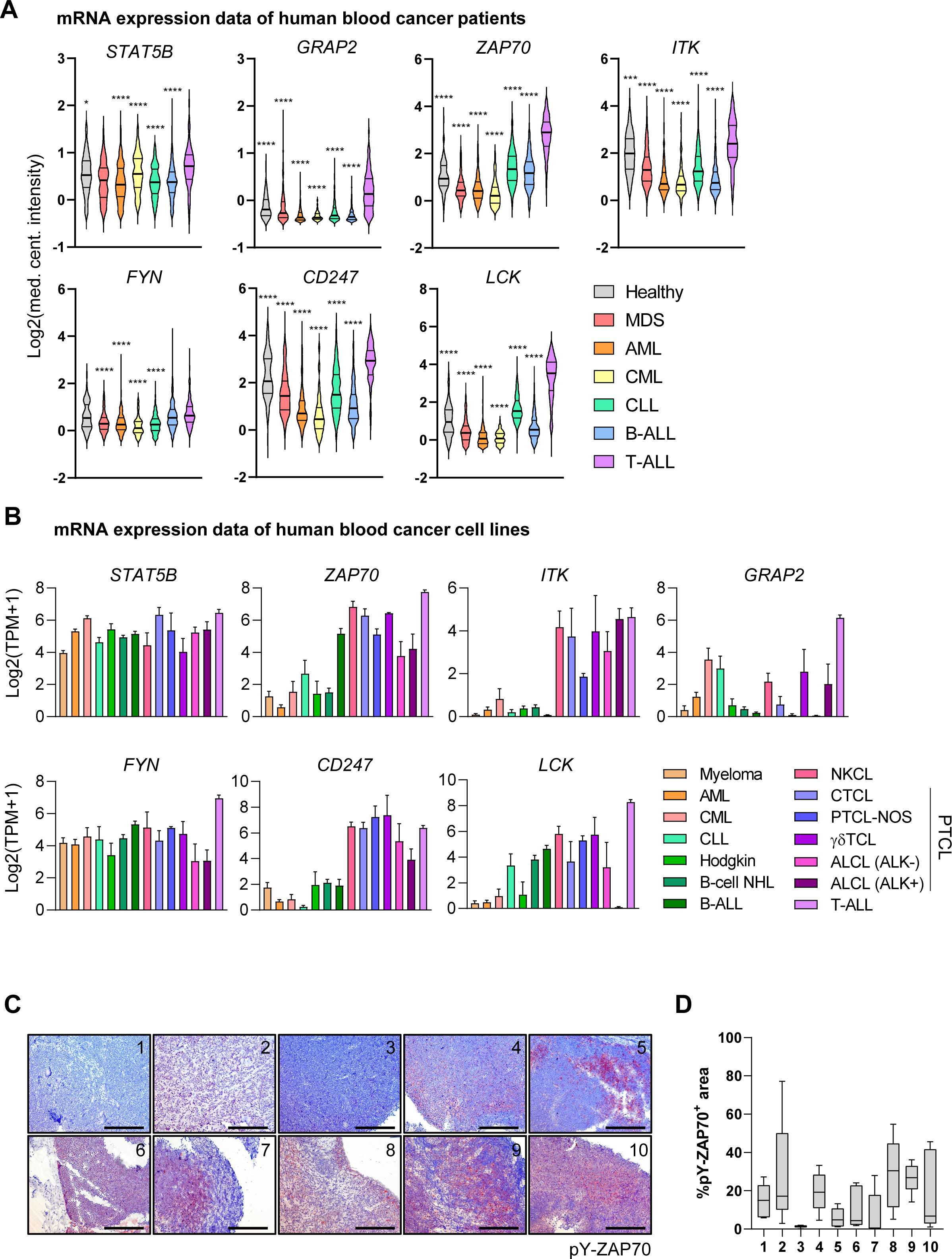
T-cell kinases are upregulated and active in human T-ALL. **A**. *STAT5B*, *GRAP2*, *ZAP70*, *ITK*, *FYN*, *CD247* and *LCK* mRNA expression data of human patients suffering from different blood cancers and healthy bone marrow control cells. Data extracted from the Haferlach Leukemia study of the Oncomine database. **B**, Expression of the same genes indicated in **A** in human blood cancer cell lines. Data extracted from DepMap (www.depmap.org). TPM: transcripts per million. Abbreviations used in **A** and **B**: MDS: myelodysplastic syndrome, AML: acute myeloid leukemia, CML: chronic myelogenous leukemia, CLL: chronic lymphocytic leukemia, B-ALL: B-cell acute lymphoblastic leukemia, NHL: non-Hodgkin lymphoma, NKCL: natural killer cell lymphoma, PTCL-NOS: peripheral T-cell lymphoma, not otherwise specified, ALCL: anaplastic large cell lymphoma. **C**, Representative immunohistochemical stainings of 10 T-ALL patient tumor samples for pY-ZAP70, original magnification: 20x, scale bar = 200 µm. **D**, Quantification of areas positive for pY-ZAP70 in patient samples shown in **C**, 6 distinct areas per slide were subjected to analysis using ImageJ and the Colour Deconvolution2 plugin. In **A**, significant differences are shown for comparisons of each dataset to T-ALL indicated as \**P* < 0.05, \*\**P* < 0.01, \*\*\**P* < 0.001, \*\*\*\**P* < 0.0001 by one-way Anova with Dunnet’s multiple comparison test. Error bars show mean +/-SEM.

### Human T-ALL cell lines with active STAT5 respond to pharmacologic inhibition of STAT3/5 or ZAP70

To assess the potential of targeting STAT5 and downstream kinases in T-ALL, we profiled nine human T-ALL cell lines for their STAT5 activity. Five T-ALL cell lines (KOPT-K1, DND-41, HSB-2, SUP-T13 and ALL-SIL) displayed high STAT5 activation by Western blotting. STAT5 activation was lower in the PEER cell line and the lowest in LOUCY, JURKAT and MOLT-4 cells, similar to the AML lines MOLM-13 and MV4-11 (**Fig. 5A**). Sanger sequencing on mRNA encoding the SH2 domain of STAT5B identified a novel, not yet described biallelic N642H driver mutation in KOPT-K1 cells (**Fig. 5B**). This explains the high level of STAT5 activation observed in this cell line and validates it as a relevant human model to study STAT5B^N642H^ in T-ALL. The DND-41 cell line lacked *STAT5B* mutations (**Fig. 5B**) but is known to carry activating *IL7R* (*p.L242_L243insLSRC*) and loss-of-function *PTPRC* (W764*) mutations, leading to active IL-7Rα-JAK1-STAT5B signaling (8) (**Fig. 6A**). None of the other tested T-ALL cell lines harbored GOF SH2 domain mutations in *STAT5B*, nor in the JH2 domain of *JAK1* or *JAK3* (further targeted Sanger sequencing data not shown).

**Figure 5.**
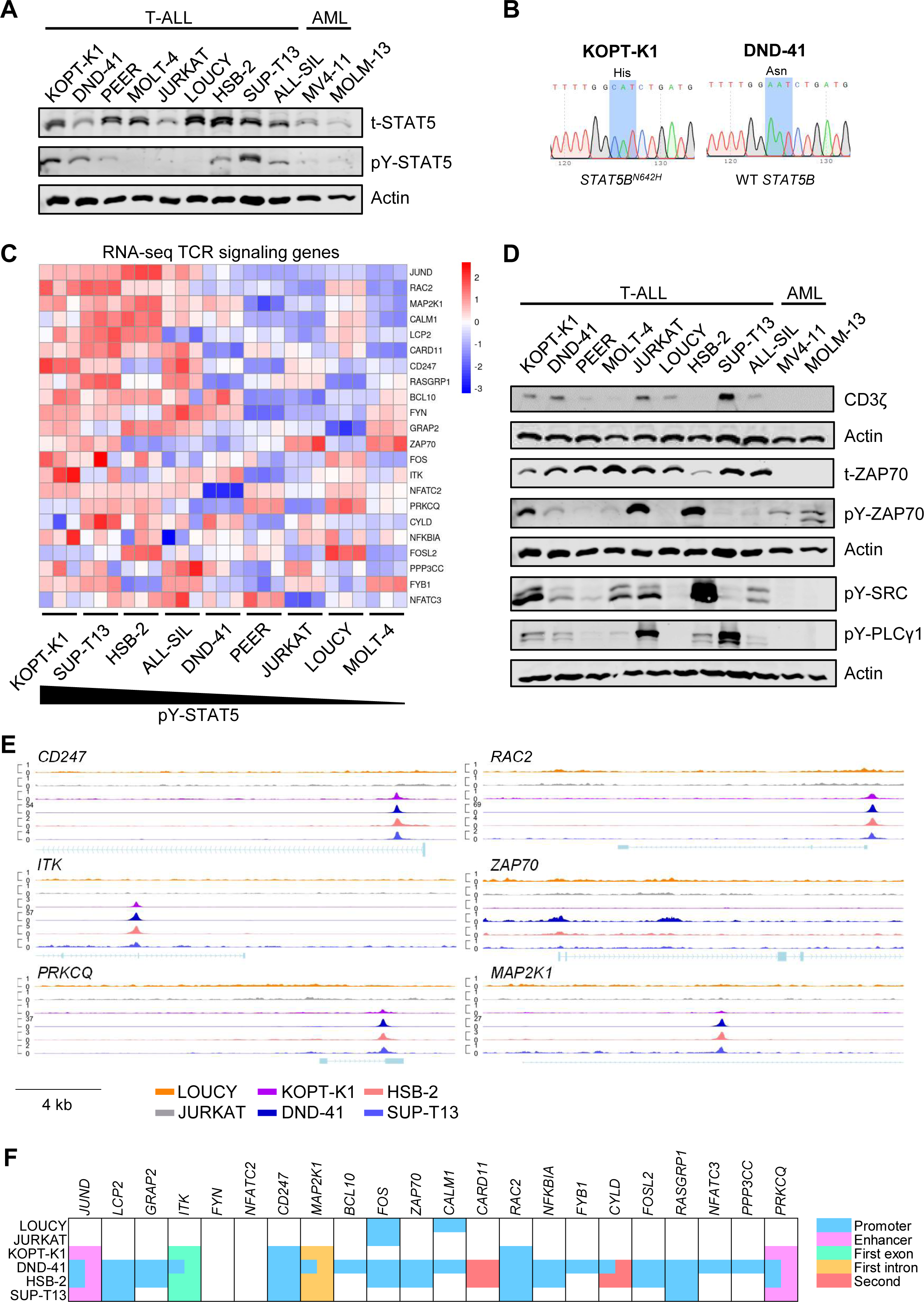
Human T-ALL cell lines with high STAT5 activation display upregulated expression of TCR pathway genes. **A**, Basal levels of pY-STAT5 and total (t) STAT5 in nine T-ALL and two AML control cell lines, evaluated by Western blot analysis, Actin served as loading control. **B**, Sanger sequencing of the SH2- and TAD-domains of STAT5B. cDNA was obtained from isolated RNA of KOPT-K1 and DND-41 cells, the codon for amino acid position 642 is highlighted in blue. **C**, Z-scores of raw counts of TCR pathway genes from RNA-seq data from nine T-ALL cell lines. Three biological triplicates were acquired from each cell line and cell lines were arranged according to their pY-STAT5 level determined by Western blot. **D**, Western blot analysis evaluating expression of CD3ζ, ZAP70 and activation of ZAP70, SRC and PLCγ1 in nine T-ALL and two AML cell lines. Actin was used as loading control. **E**, ChIP-seq signals indicating STAT5B binding at the promoter region of 6 TCR pathway genes as indicated in 6 T-ALL cell lines, normalized ChIP-seq signal (CPM) indicated on the Y-axis. **F**, Peak analysis for STAT5B binding sites in TCR pathway genes from six T-ALL cell lines and indication of peak localization within respective genes or enhancers.

**Figure 6.**
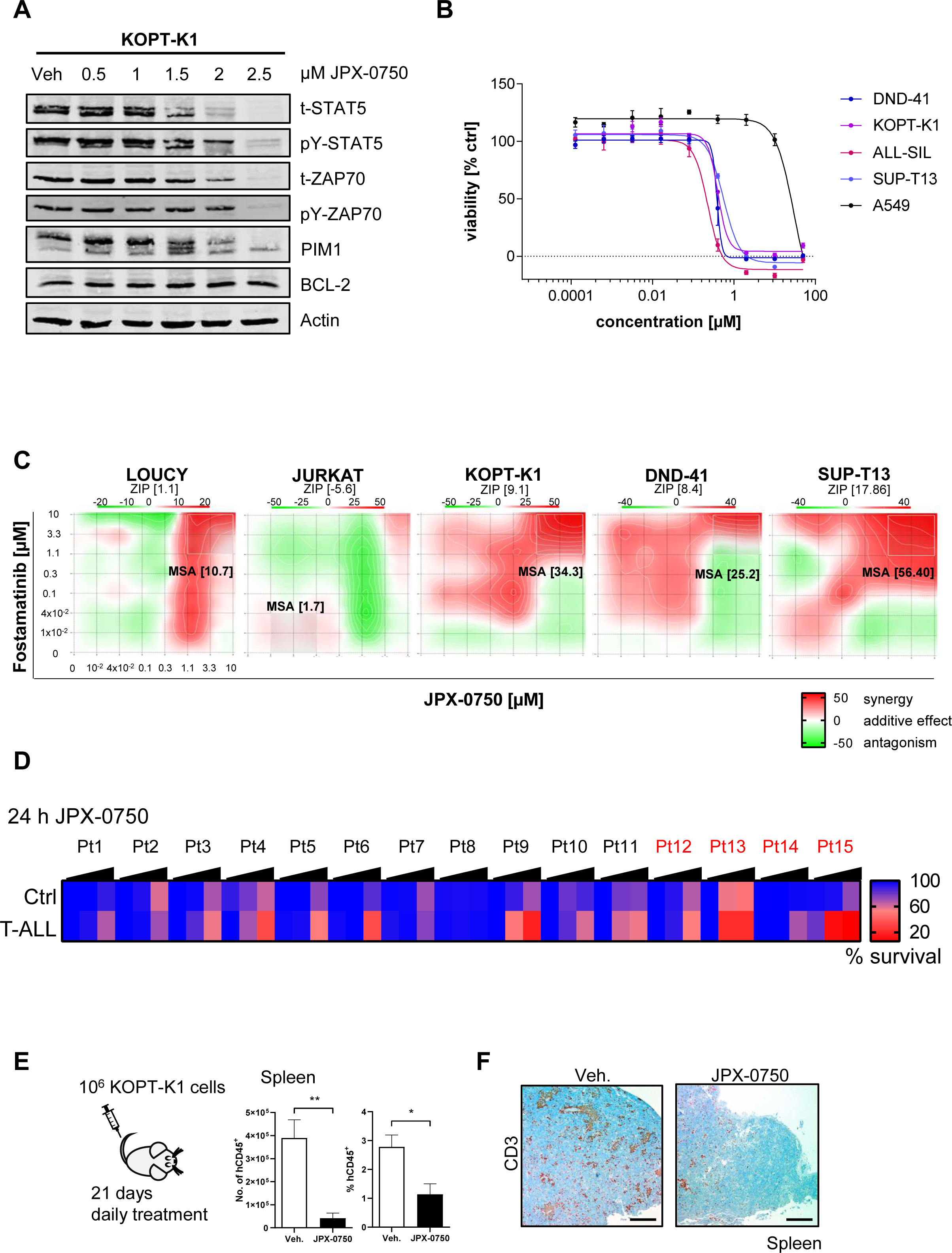
T-ALL cells with elevated TCR pathway gene expression and STAT5 activation respond to ZAP70 and STAT5 inhibition. **A**, Western blot analysis for pY-STAT5, t-STAT5, PIM1 and BCL-2 of T-ALL cells after treatment with STAT5 degrader JPX-0750 at indicated concentrations or vehicle (Veh.) for 24h. α-Tubulin was used as loading control. **B**, Dose-response curve for T-ALL cell lines treate with JPX-0750 at indicated concentrations determined using Cell-TiterBlue viability assay. Three independent experiments in technical triplicates were performed. **C**, Dose-response synergy analysis of the indicated two-drug combination in T-ALL cells after 72 h treatment. In each graph, the most synergistic area (MSA) is highlighted (white rectangle), which represents the most synergistic 3-by-3 dose-window with the respective MSA score. One representative plot out of two independent experiments is shown. **D**, Percent survival of healthy control and T-ALL cells of primary patient samples treated 24 h with JPX-0750 at 100, 1000 or 10,000 nM. Mean values of respective concentration vs. DMSO control in 2 technical replicates are shown. Relapsed patients are marked in red. **E**, KOPT-K1 cell numbers in the spleen after transplantation into NSG mice, and treatment with JPX-0750 (n = 5) or vehicle (n = 4) for 21 days. **F**, Representative immunohistochemical stainings for CD3 in the spleens of KOPT-K1 transplanted NSG mice, treated with vehicle or JPX-0750, original magnification: 4x, scale bar = 500 µm. In **F**, significant differences are indicated as

Next, we performed RNA-seq of T-ALL cell lines to evaluate the impact of STAT5 activity on gene transcription of TCR pathway members further. We found higher expression of these genes in cell lines with higher STAT5 activation (**Fig. 5C**). In line, we found higher CD3ζ chain and ZAP70 protein expression, where ZAP70 and downstream SRC and PLCγ1 activation in T-ALL cell lines confirmed active kinase signaling downstream of the TCR in T-ALL (**Fig. 5D**).

To further elucidate whether STAT5 is directly involved in TCR pathway gene regulation, we performed ChIP-seq analysis for STAT5B in four T-ALL cell lines with high pY-STAT5 levels (KOPT-K1, DND-41, HSB-2 and SUP-T13) and two with low pY-STAT5 levels (LOUCY and JURKAT, **Fig. 5A**). The robustness of our ChIP-seq analysis was confirmed by STAT5B peaks in the *bona fide* STAT5 target genes *BCL2*, *BCL2L1*, *SOCS3* and *RUNX3* in KOPT-K1, DND-41, HSB-2 and SUP-T13 cells in (**Supplementary Fig. S11A**). Furthermore, we found that STAT5 binds at promoters, enhancers, introns or exons of TCR pathway genes including *CD247* and downstream effector molecules *ZAP70*, *ITK*, *MAP2K1*, *PRKCQ*, *RAC2* and others (**Fig. 6E** and **F**), specifically in the cell lines with high STAT5 activity. This indicates that these genes could be transcriptional targets upregulated by active STAT5 in particular.

We therefore reasoned that T-ALL cells might be sensitive to pharmacologic inhibition of STAT5 or the important T-cell kinase ZAP70. We tested this using JPX-0750 (52,53), a novel, small molecule STAT3/5 degrader that was developed to prevent compensatory STAT3 upregulation upon STAT5 targeting as an escape route. Thus, JPX-0750 could be a suitable lead compound available to degrade oncogenic STAT3/5, or the established SYK/JAK inhibitors Fostamatinib or Gusacitinib, since ZAP70 is one of two members of the SYK protein family. Gusacitinib is a dual SYK/JAK inhibitor for which the FDA has granted fast track designation for chronic hand eczema (54). Fostamatinib is a SYK inhibitor FDA-approved for chronic immune thrombocytopenia (55). Treatment of T-ALL cells with JPX-0750, rapidly abrogated STAT5 and ZAP70 activation and expression, suggesting direct dependency on STAT3/5. JPX-0750 treatment induced diminished levels of PIM1, but BCL-2 expression remained normal (**Fig. 6A**). While *PIM1* and *BCL2* are STAT5 target genes, BCL-2 is also independently regulated by the PI3K/AKT/mTOR pathway, which is in turn interwoven with STAT5 signaling in T-cell metabolism (45,56). T-ALL cells with high STAT5 activation were sensitive to JPX-0750 compared to control lung adenocarcinoma A549 cells (**Fig. 6B**). Moreover, we found remarkable responses of T-ALL cells with high pY-STAT5 levels and kinase activation to Fostamatinib or Gusacitinib treatment. Fostamatinib displayed a higher selectivity to those cell lines with higher kinase activation or ZAP70 expression (KOPT-K1, HSB-2, JURKAT, ALL-SIL, DND-41, SUP-T13, MOLT-4, **Supplementary Fig. S11B**). Importantly, T-ALL cells treated with these selective kinase blockers displayed reduced levels of ZAP70 and STAT5 activation without significantly altering their total protein levels (**Supplementary Fig. S11C**). JAK/STAT and pre-TCR signaling are distinct pathways and our findings suggest cooperative cross-regulation. Thus, we tested whether combined inhibition leads to synergy. Notably, Fostamatinib synergized with JPX-0750 selectively in KOPT-K1, DND-41 and SUP-T13 cells with high pY-STAT5 and elevated TCR pathway gene expression, while no significant synergy was observed in the control cell lines LOUCY and JURKAT (**Fig. 6C**). Hence, our data suggest that combinatorial treatment with SYK and STAT5 inhibitors appear promising for T-ALL patients with activated STAT5/ZAP70. Next, we treated primary human T-ALL cells (**Supplementary Table 3**) with JPX-0750 or Fostamatinib and found higher sensitivity to JPX-0750 in T-ALL cells compared to non-tumor cells in 12 out of 15 samples (**Fig. 6D**). Fostamatinib selectively killed T-ALL cells in two samples, importantly, the sample that responded best to JPX-0750 (Patient 10) was also highly sensitive to Fostamatinib (**Supplementary Fig. S11D**). Remarkably, both Fostamatinib-responders were relapsed T-ALL patients, indicating that more progressive and aggressive disease stages might be more sensitive to ZAP70 inhibition (**Supplementary Table 3**). Since JPX-0750 revealed the best responses *in vitro*, we used this inhibitor to confirm its efficacy *in vivo*. Strikingly, treatment with JPX-0750 resulted in a reduced number of intravenously transplanted KOPT-K1 cells in the spleens of immunocompromised mice (**Fig. 6E** and **6F**). Altogether, we conclude that mutated STAT5A and STAT5B are oncoproteins in T-ALL. Their hyperactivation drives upregulation of genes of the CD3ζ-ZAP70 axis, representing future vulnerable signaling nodes for targeted therapy.

## Discussion

We establish that STAT5A and STAT5B GOF oncoproteins reprogram thymic T-cell development and promote leukemic transformation of immature T-cells, giving rise to T-ALL. We suggest that hyperactive STAT5 mimics pre-TCR signaling in T-ALL, by directly activating transcription of TCR pathway genes. We discovered a biallelic *STAT5B^N642H^* mutation in the KOPT-K1 cell line defining it as a patient-derived model to study the most frequent STAT5B driver mutation. Furthermore, we validated that STAT5 and ZAP70 signaling are essential in a subset of human T-ALL. Pharmacologic intervention using STAT5/SYK inhibitors either alone or synergistically blocked human T-ALL cell growth and survival. Due to the significant rate of *STAT5B* mutations in nine different T-cell cancers, exhibiting STAT5 activation status and the essential genetic role for STAT5 in T-cell differentiation, function, proliferation and survival, we postulate, that these findings extend from T-ALL also to mature T-cell cancers such as PTCL, which comprise a large group of ∼30 different T-cell cancers (57). This is supported by the fact that activating *STAT5B* mutations and particularly *STAT5B^N642H^* also occur in T-cell large granular lymphocytic leukemia, T-cell prolymphocytic leukemia and various γδ T-cell derived leukemia/lymphoma (19). Furthermore, STAT5 activation is targetable in cutaneous T-cell lymphomas (CTCL), bearing *STAT5A*/*B* copy-number-gains and PDGFRβ-driven anaplastic large cell lymphoma (53,58). STAT5 activating kinase triggers might lead to similar molecular consequences as *STAT5B^N642H^*, highlighting broad applicability of our results. STAT5A/B have been validated as oncogenic drivers of T-cell malignancies in mouse models (17–19). Higher basal expression levels of STAT5B over STAT5A likely contribute to a dominant role of STAT5B in T-cell biology and leukemogenesis and hyperactive STAT5A requires stronger overexpression to transform T-cells (18). This might explain the lower impact of STAT5A^S710F^ on DN1-4, DP, SP4/8 stage composition compared to STAT5B^N642H^. Comprehensive RNA-seq analysis revealed enhanced T-cell activation and lineage commitment with significant RUNX3, EOMES or T-bet upregulation by STAT5B^N642H^ or STAT5A^S710F^ expression in purified DN, DP and SP8 thymocytes. Our data display parallels with findings from the ÓShea and Hennighausen groups, demonstrating that murine *Stat5b* knockout had a higher impact on T-cell differentiation and function than *Stat5a* knockout, whereas individual *Stat5a* or *Stat5b* knockouts were highly similar at the transcriptome level (59). Normally, T-cell development and selection are highly sensitive processes, where physiological negative and positive selection lead to elimination of up to 95% of thymocytes (60). Thus, it is surprising that T-cells bearing hyperactive STAT5 most-likely overcome apoptosis in thymic selection processes in the thymus and give rise to mature, post-thymic T-cells (17).

Studies identified mutated or overexpressed IL-7R, JAK1 or JAK3 as oncoproteins in T-ALL patients, and validated them as oncogenic drivers in mouse models (8,9,12,14,61). Many of these studies suggested IL-7R-JAK1/3 signaling via STAT5. The question whether *STAT5A* or *STAT5B* GOF mutations can directly induce T-ALL remained unanswered. STAT5 GOF alone failed to induce a T-ALL-characteristic arrest in T-cell development in the presence of VDJ recombination. To counteract mature T-cell generation, we induced arrest in thymic development in STAT5 GOF mice by crossing them with RAG2^-/-^ mice. Indeed, these mice succumbed to a T-ALL-like disease, mainly characterized by massive thymic enlargement and dominant thymic TdT expression. RAG2-deficiency may imitate other common genetic lesions in human T-ALL, such as chromosomal rearrangements (4). Whether STAT5 mutations cooperate with these lesions, e.g. involving *TAL1*, *HOXA1* or *TLX1* or with hyperactive *NOTCH1* mutations remains to be explored. We reveal that both STAT5 GOF variants launch a transcriptional program that is transcriptionally similar to ETP-ALL. However, the lack of comprehensive, subtype-specific transcriptomes of human patients exacerbates definite conclusions about the T-ALL subtype of our mice. Moreover, STAT5B^N642H^ occurs in several clinical T-ALL subtypes (4,24), highlighting the genetic heterogeneity of this disease. Most tumor cells in STAT5 GOF RAG2^-/-^ mice are DP or SP8, suggesting progression in thymic development beyond the DN stage, despite TCR-deficiency due to loss of VDJ rearrangement. This resembles induction of thymocyte differentiation promoted by LCK- or NPM-ALK-overexpression or CD3 cross-linking in RAG-deficient mice (46,62). However, comprehensive analyses to better understand the transcriptional regulation of these processes by NGS efforts have not been performed. Our RNA-seq data provide mechanistic explanations revealing a pronounced transcriptional upregulation of genes encoding kinases or adapter proteins that operate downstream of the TCR pathway.

Translating our findings, we found that high expression of these genes is specific to T-ALL as compared to other more mature T-cell cancer types in humans. Most interestingly, their expression values are higher in T-ALL than in PTCL, where aberrant TCR signaling and its targeting has been proposed in several disease subtypes (48–51). Our findings in mice are consistent with a transcriptional upregulation of important TCR signaling genes such as *CD247*, *JUND*, *RAC2*, *MAP2K1*, *CALM1*, *LCP2*, *FYN*, *GRAP2* and *ZAP70,* correlating with STAT5 activity in human T-ALL cells. Moreover, ChIP-seq analysis revealed that STAT5B binds to the promoter of the *CD247* gene. CD3ζ transduces signals to the SYK family member ZAP70 and thereby activates TCR-mediated T-cell responses. We observed a weaker upregulation of TCR pathway genes by hyperactive STAT5A in the absence of RAG2 expression, with delayed T-cell neoplasia development compared to more aggressive STAT5B^N642H^-driven disease. Protein crystallization studies implicated that STAT5B^N642H^ mediates structural changes that lead to fortified dimerization accompanied with delayed dephosphorylation adjacent to the SH2-pY interaction and thereby sustained STAT5 activity. Moreover, STAT5B^N642H^ can both render cells cytokine-independent or hypersensitive to upstream cytokine stimulation (19). This is consistent with recent findings of the Cools laboratory, showing that cells expressing STAT5B^N642H^ are sensitive to JAK inhibitors, indicating dependency on upstream kinase nodes (63). This concept challenges the classic understanding of pathways to signal in one direction. Whether T-ALLs with STAT5 GOF mutations are still cytokine- or JAK-dependent and if similar crosstalks operate between STAT5 and other kinases, such as ZAP70, ITK or FYN, remains to be explored.

Previous studies revealed that T-ALL cells are sensitive to Dasatinib, targeting the kinase activity of the TCR pathway member LCK (64,65). It was demonstrated through shRNA knockdowns of six TCR pathway members as well as Dasatinib treatment that T-ALL cell lines and patient derived xenografts (PDX) are LCK-dependent. Moreover, Dasatinib synergized with dexamethasone treatment, suggesting combination therapy including pharmaceutical targeting of T-cell kinases as a promising treatment option (66). It had also been previously shown that three T-ALL cell lines were sensitive to ZAP70 knockdown, but the molecular mechanisms of ZAP70 upregulation remained unknown and targeting of ZAP70 has not been tested in T-ALL (66). We found that the STAT5 inhibitor JPX-0750 or the ZAP70 tyrosine kinase inhibitors Fostamatinib and Gusacitinib effectively blocked T-ALL cell line growth and survival. Importantly, the combination of STAT5 and ZAP70 inhibitors acted synergistically in T-ALL cell line growth/survival inhibition, arguing that these pathways act independently to regulate survival and growth in T-ALL. JPX-0750 specifically killed T-ALL cells in the vast majority of primary human T-ALL samples, but a lead candidate as a small molecular weight STAT3/5 degrader moving into a clinical trial does not exist yet. Moreover, it remains unclear, why Fostamatinib affected only 2 out of 15 samples. However, higher numbers of samples for testing might be required due to high heterogeneity between T-ALL subtypes and other kinases might also be involved. In summary, our data reveal how *STAT5A* and *STAT5B* GOF mutations affect thymic T-cell development and immature T-cell transformation, leading to T-ALL. We uncover novel mechanisms showing that hyperactive STAT5 can directly mimic pre-TCR pathway action and we pioneer the principle of STAT5 or SYK kinase family targeting. We conclude that kinase or direct STAT5 inhibition are promising strategies to block T-ALL growth, but further medicinal chemistry programs and clinical work is needed to move targeted drugs to patients.

## Materials and Methods

### Mice

All mice used were on C57BL/6, C57BL/6-Ly5.1 (B6.SJL-*Ptprc^a^Pepc^b^*/BoyCrl) or NSG (NOD.Cg-*Prkdc^scid^ Il2rg^tm1Wjl^*/SzJ) background and maintained in a specific-pathogen–free quality in the experimental mouse facility at the University of Veterinary Medicine (Vienna, Austria). Transgenic mice hemizygous for human *STAT5B^N642H^*, human WT *STAT5B* or murine *Stat5a*^S710F^ driven by the *Vav1*-promoter (67) and 3’ *Vav1*-enhancer were generated and genotyped as previously described (17,18). Double mutants were produced by cross-breeding the three transgenic lines with *Rag2^tm1Fwa^* mice and subsequent breeding for homozygosity of the *Rag2* knockout allele. Due to rapid disease development STAT5B^N642H^ mice were also generated via *in vitro* fertilization with frozen sperm cells.

### Flow cytometry

For flow cytometry, single-cell suspensions were prepared by crushing organs through a 70 μm cell strainer (BD Biosciences). Erythrocytes of spleens were lysed using Ammonium-Chloride-Potassium (ACK) buffer (150 mM NH_4_CO_3_, 10 mM KHCO_3_, 1 mM EDTA, pH 7.2). All antibodies used for flow cytometry are listed in Supplementary Methods Table 1. All analyses were performed on the BD FACSCanto II using FACSDiva (BD) and FlowJo (version 10.5.3) software. Sorting was performed on a BD FACSAria II.

### Blood analysis

Mice were anesthetized and blood was drawn via heart puncture and transferred into EDTA tubes. WBC counts were determined using the scil VetABC device.

### RNA-seq sample acquisition

DN, DP and SP8 T-cells were harvested from thymi by generating single-cell suspensions and FACS-sorting for viable Thy1.2-positive cells with respective CD4/CD8 expression patterns was performed using fluorescent antibodies listed in the Supplementary Methods Table 1. RNA was isolated from snap-frozen cell pellets using the DNA/RNA Allprep Kit (QIAGEN). A detailed protocol of RNA-seq processing and analysis is described in the Supplementary Methods.

### Cell culture

KOPT-K1 cells were kindly provided by Koshi Akahane, from the University of Yamanashi. All other T-ALL and cell lines were kindly provided by Takaomi Sanda, from the Cancer Science Institute of Singapore. AML and A549 control cell lines were purchased from DMSZ.

All cells were cultivated with complete RPMI 1640 or DMEM medium (10% FCS, 2 mM L-glutamine, 10 U/mL penicillin-streptomycin), all from Gibco, Thermo Fisher Scientific. The cell lines were authenticated and regularly tested for *Mycoplasma* using the MycoAlert PLUS detection kit by Lonza (LT07-710).

### Western blot

Immunoblotting was performed using standard protocols and antibodies as listed in Supplementary Methods Table 2. Images of membranes were obtained using IRDye fluorescent secondary antibodies and an Odyssey CLx imaging system (LI-COR). Signal quantification was performed using Image Studio Lite (version 5.2).

### Immunohistochemistry

Mouse organs were processed and analysed using standard protocols, details and antibody specifications are described in the Supplementary Methods.

### Transplantation experiments

STAT5B^N642H^ mice were sacrificed and 10^6^ viable CD3^+^ CD8^+^ lymphocytes (from LNs) or Thy1.2^+^ CD4^+^ CD8^+^ DP thymocytes (from thymus) were isolated by flow cytometry and transplanted into Ly5.1 or C57BL/6Nrj recipients via tail vein injection. Recipients were monitored for signs of disease daily and sacrificed at indicated time points or at a moribund state.

### Human database analyses

Human patient gene expression data were extracted and downloaded from the Haferlach Leukemia, Zhang Leukemia or Andersson Leukemia datasets using the Oncomine™ Research Premium Edition database (68). Human cell line gene expression data were downloaded from DepMap (www.depmap.org).

### Human patient analysis

DNA and RNA was isolated from total leukocytes of primary T-ALL patient samples, and whole-genome-sequencing and RNA-seq was performed as described previously (69).

### Sanger sequencing

Cells were harvested, RNA was isolated using the AllPrep DNA/RNA Mini Kit from QIAGEN and cDNA was generated using the RevertAid First Strand cDNA Synthesis Kit from ThermoFisher Scientific. A sequence spanning over the mRNA region encoding the STAT5B C-terminal region was amplified by PCR (forward primer: GGCAATGGTTTGACGGTG, reverse primer: GGATCCACTGACTGTCCATT). The 646 bp PCR product was separated on an agarose gel using standard conditions. DNA was isolated using the MinElute PCR Purification Kit and Sanger sequencing was performed by Microsynth Austria GmbH (primer: GCCTCATTGGAATGATGG).

### ChIP-seq and initial processing

ChIP-seq was performed as previously described (70). The antibodies used for each experiment are listed in the **Supplementary Methods Table 3**. For each ChIP, 5 μg of antibody coupled to 2 μg of magnetic Dynabeads (Life Technologies) was added to 3 mL of sonicated nuclear extract from formaldehyde-fixed cells. Chromatin was immunoprecipitated overnight, cross-links were reversed, and DNA was purified by precipitation with phenol:chloroform:isoamyl alcohol. DNA pellets were resuspended in 25 μL of TE buffer. Illumina sequencing, library construction, and ChIP-seq analysis methods were previously described. The ChIP-seq analysis is further described in the **Supplementary Methods**.

### *In vitro* drug treatments

10^4^ cells/well were seeded in triplicates in a flat-bottom 96-well plate. On the subsequent day, cells were treated with serial 1:2 dilutions of the drug of interest. Bortezomib (MedChemExpress, HY-10227), a proteasome blocker, served as positive control and DMSO as negative control. DMSO concentrations were equal in all wells to exclude effects by variations in concentrations of the vehicle. Treated cells were incubated at 37°C and 5% CO_2_ for 72 h. Cell viability was assessed by using the CellTiter-Glo LuminescenT-cell Viability Assay (Promega) or the CellTiter-Blue Cell Viability Assay (Promega) and the GloMax® Discover Microplate Reader (Promega). IC_50_ values were calculated from ATP luminescence using non-linear regression of log-transformed, normalized data.

For Western blot analysis, 1.5x10^6^ cells were seeded in 2 mL medium per well in 6-well plates. On the next day, cells were treated with indicated doses of JPX-0750, Fostamatinib (MedChemExpress, HY-12067) or Gusacitinib (MedChemExpress, HY-103018). After 24 h, cells were harvested into 15 mL tubes, washed with ice-cold PBS and processed for Western blot analysis.

### Drug synergy testing

The drug combination synergy analysis was performed using the SynergyFinder 2.0 web application. The degree of synergy was quantified using the zero interaction potency (ZIP) model, which captures the drug interaction relationships by comparing the change in the potency of the dose–response curves between individual drugs and their combinations. In addition to the overall ZIP score over the dose-response matrix, the SynergyFinder web-tool enabled calculation of the most synergistic area (MSA) score, which represents the most synergistic 3-by-3 dose-window. According to the ZIP model, an MSA score below -10 indicates that the two drugs have antagonistic effects, a score between -10 and 10 indicates an additive effect, while a MSA score above 10 indicates a synergistic effect. The four-parameter logistic regression (LL4) model was used as the curve-fitting algorithm in the SynergyFinder web-tool. All assays were performed in two biological replicates.

### Chemosensitivity profiling in primary patient cells

T-ALL patient samples of bone marrow aspiration or peripheral blood were obtained from the Division of Hematology and Hemostaseology at the Medical University of Vienna (Vienna, Austria). Each sample was frozen after Ficoll-Paque mononuclear cell separation. Fostamatinib and JPX-0750 were printed on tissue culture-treated 384-well plates (Corning) using an Echo 550 (Labcyte Inc.) acoustic dispenser in five different concentrations in 10-fold dilutions encompassing a 10,000-fold concentration range (1 - 10,000 nM) in duplicates (only 100, 1000 and 10,000 nM responses were taken into analysis). DMSO was used as a negative control. Samples were thawed, resuspended in RPMI + 10% FBS, and seeded at a concentration of 4-5x10^5^ per mL (50 mL/well) onto pre-drugged 384-well plates. The cells with drugs were incubated at 37°C for 24 h. After incubation, the cells were spun down (100 g for 5 min) and the supernatant from each well was aspirated using the Biotek MultiFlo FX Multi-Mode Dispenser. The cells were stained with DAPI and antibodies marking T-ALL or control cells (Supplementary Methods Table 3) at a 1:500 dilution in PBS for 1 h at room temperature. Then, the plates were run on an iQue3 high-throughput flow cytometer screener (Sartorius), which facilitated multiplex cell analysis and subclone specific drug screening. The data was gated to remove noise, doublets, and dead cells. T-ALL cells of each sample were determined evaluating histological reports and FACS marker positivity.

### Xenograft experiment

10^6^ KOPT-K1 cells were transplanted via tail vein injection into NSG recipients. Transplanted cells were traced by regular bleedings and subsequent flow cytometry staining for surface human CD45 to verify engraftment in all mice used for the treatment. Recipients were randomized and treated daily by intraperitoneal injection with JPX-0750 at a dosage of 10 mg/kg or vehicle for 21 subsequent days, starting 9 days after injection. JPX-0750 was dissolved in DMSO and diluted 1:100 in 50% PEG-400, 5% TWEEN®80, 44% PBS.

### Ethics approval

All mouse experiments were approved by the institutional ethics committee of the University of Veterinary Medicine Vienna and licensed under BMWF-68.205/0023-II/3b/2014, BMWFW-68.205/0166-WF/V/3b/2015, BMBWF-68.205/0130-WF/V/3b/2016 and BMBWF-68.205/0041-V/3b/2019 by the Austrian Federal Ministry of Education, Science and Research.

Material from human patients was obtained with approval from the ethics committees of the Medical University of Vienna (vote number EK 1727/2022). Research was conducted in accordance with the Declaration of Helsinki.

Tissue samples used for immunohistochemistry were provided by the University Cancer Center Frankfurt (UCT). Written informed consent was obtained from all patients in accordance with the Declaration of Helsinki and the study was approved by the institutional review boards of the UCT and the Ethics Committee at the University Hospital Frankfurt (project-number: SHN-06-2018).

## Availability of data and materials

All data that were used for analyses in this study can be requested from the corresponding author. RNA-seq and ChIP-seq data were uploaded to the GEO database repository and can be downloaded via the accession numbers GSE218858 and GSE219155 after permission from the corresponding author.

## Authors’ contributions

RM designed and supervised the study. TS, HS, NP, MWZ, BM, SS, KS, AP, TP, CS, DP, DJ, TAM, MMKA, HTTP and KZ designed, performed, and/or analyzed experiments. FR, MWZ and TE contributed bioinformatics support. MD, AB and TR generated and maintained transgenic mice. TK, CB, MM, RF, PTG, TA, SM, TSa, SH, FG, GH, TH, PBS, HAN, ATL and MH contributed to the interpretation of data and revised the manuscript. RM and TS wrote the manuscript.

## Acknowledgments

We would like to thank Safia Zahma and Michael Machtinger from the Institute of Animal Breeding and Genetics at the University of Veterinary Medicine Vienna, Vienna, for their support in histology and mouse handling and processing. Moreover, we highly appreciate the computational support by Brian J. Abraham from the Division of Molecular Oncology, St. Jude Children’s Research Hospital, Memphis, TN and Stephan Hutter and Wecke Walter from the Munich Leukemia Laboratory, Munich.

## List of abbreviations

AGO2: Argonaute protein 2
AML: Acute myeloid leukemia
BCL-2: B-cell lymphoma 2
(c)DNA: (complementary) deoxyribonucleic acid
ChIP: Chromatin immunoprecipitation
CRISPR: Clustered Regulatory Interspaced Short Palindromic Repeats
CTCL: Cutaneous T-cell lymphoma
DMSZ: Deutsche Sammlung von Mikroorganismen und Zellkulturen
DN: Double-negative
DP: Double-positive
EOMES: Eomesodermin
ETP: Early T-cell progenitor
FACS: Fluorescence-activated cell sorting
fc: Fold change
FDA: Food and drug administration
FOXP3: Forkhead box P3
GOF: Gain-of-function
GRAP2: GRB2-related adapter protein 2
GSEA: Gene set enrichtment analysis
H&E: Hematoxylin and eosin
HOXA: Homeobox protein A
HOXA: Homeobox protein A
IL7R: Interleukin 7 receptor
ITK: Interleukin-2-inducible T-cell kinase
JAK: Janus kinase
JH2: JAK homology domain 2
LCK: Lymphocyte-specific protein tyrosine kinase
LMO1/2: Rhombotin-1/2
LN: Lymph node
MHC: Major histocompatibility complex
MSA: Most synergistic area
mTOR: Mammalian Target of Rapamycin
NK: Natural killer
NSG: NOD-SCID Il2rgnull
Padj: Adjusted P value
PDGFR: Platelet Derived Growth Factor Receptor
PDX: Patient-derived Xenograft
PI3K: Phophatidylinositol 3-kinase
PIM1: Proto-oncogene serine/threonine-protein kinase Pim-1
PTCL: Peripheral T-cell lymphoma/leukemia
pY: Phospho-Tyrosine
RAG2: Recombination activating gene 2
RNA: Ribonucleic acid
RUNX3: Runt-related transcription factor 3
seq: Sequencing
SH2: SRC Homology 2
SP: Singe-positive
SRC: Sarcoma kinase
STAT: Signal transducer and activator of transcription
SYK: Spleen tyrosine kinase
TAL1/2: T-cell acute lymphocytic leukemia protein 1/2
T-ALL: T-cell acute lymphoblastic leukemia
T-bet: T-box transcription factor TBX21
TCR: T-cell receptor
TdT: Terminal deoxynucleotidyl transferase
T-LBL: T-cell lymphoblastic lymphoma
TLX1/3: T-cell leukemia homeobox protein 1/3
WBC: White blood cell
WT: Wild type
ZAP70: Zeta-chain-associated protein kinase 70
ZIP: Zero interaction potency

## Supplementary Tables

**Supplementary Table 1.** Correlation analysis by GSEA between RNA-seq of DN, DP and SP8 cells of STAT5B^N642H^ vs. WT and characteristic signatures of T-cell subsets, further described in reference (71). Number of genes in each gene set (SIZE), enrichment score (ES), normalized enrichment score (NES), nominal *P* value (NOM *P*), false discovery rate, q value (FDR q-val) and family-wise error rate, q value (FWER q-val) are shown.

| GS DETAILS | SIZE | ES | NES | NOM p-val | FDR q-val | FWER p-val |  |
| --- | --- | --- | --- | --- | --- | --- | --- |
| IFN RESPONSE | 50 | 0.74 | 2.69 | 0 | 0 | 0 | DN |
| CD8 CYTOTOXIC | 36 | 0.71 | 2.4 | 0 | 0 | 0 |  |
| CD8 CYTOKINE | 49 | 0.61 | 2.2 | 0 | 0 | 0 |  |
| CD4 NV/CM<br>REST | 40 | 0.54 | 1.88 | 0 | 0.001 | 0.002 |  |
| CD4/CD8 | 46 | 0.5 | 1.81 | 0.001 | 0.001 | 0.004 |  |
| TREG | 48 | 0.48 | 1.74 | 0.004 | 0.003 | 0.015 |  |
| IFN RESPONSE | 50 | 0.85 | 3.08 | 0 | 0 | 0 | DP |
| CD8 CYTOKINE | 42 | 0.81 | 2.85 | 0 | 0 | 0 |  |
| CD8 CYTOTOXIC | 28 | 0.75 | 2.42 | 0 | 0 | 0 |  |
| TREG | 48 | 0.55 | 2.01 | 0 | 0.001 | 0.002 |  |
| CD4/CD8 | 42 | 0.55 | 1.9 | 0 | 0.001 | 0.004 |  |
| CD4 NV/CM | 45 | 0.4 | 1.42 | 0.047 | 0.067 | 0.246 |  |
| PROLIFERATION | 45 | 0.31 | 1.09 | 0.345 | 0.339 | 0.849 |  |
| CD8 CYTOTOXIC | 32 | 0.78 | 2.62 | 0 | 0 | 0 | SP8 |
| CD8 CYTOKINE | 44 | 0.71 | 2.58 | 0 | 0 | 0 |  |
| IFN RESPONSE | 49 | 0.64 | 2.36 | 0 | 0 | 0 |  |
| CD4/CD8 | 45 | 0.49 | 1.79 | 0.002 | 0.003 | 0.009 |  |
| TREG | 48 | 0.38 | 1.37 | 0.066 | 0.1 | 0.304 |  |
| CD4 NV/CM | 41 | 0.3 | 1.08 | 0.343 | 0.341 | 0.827 |  |

**Supplementary Table 2.**
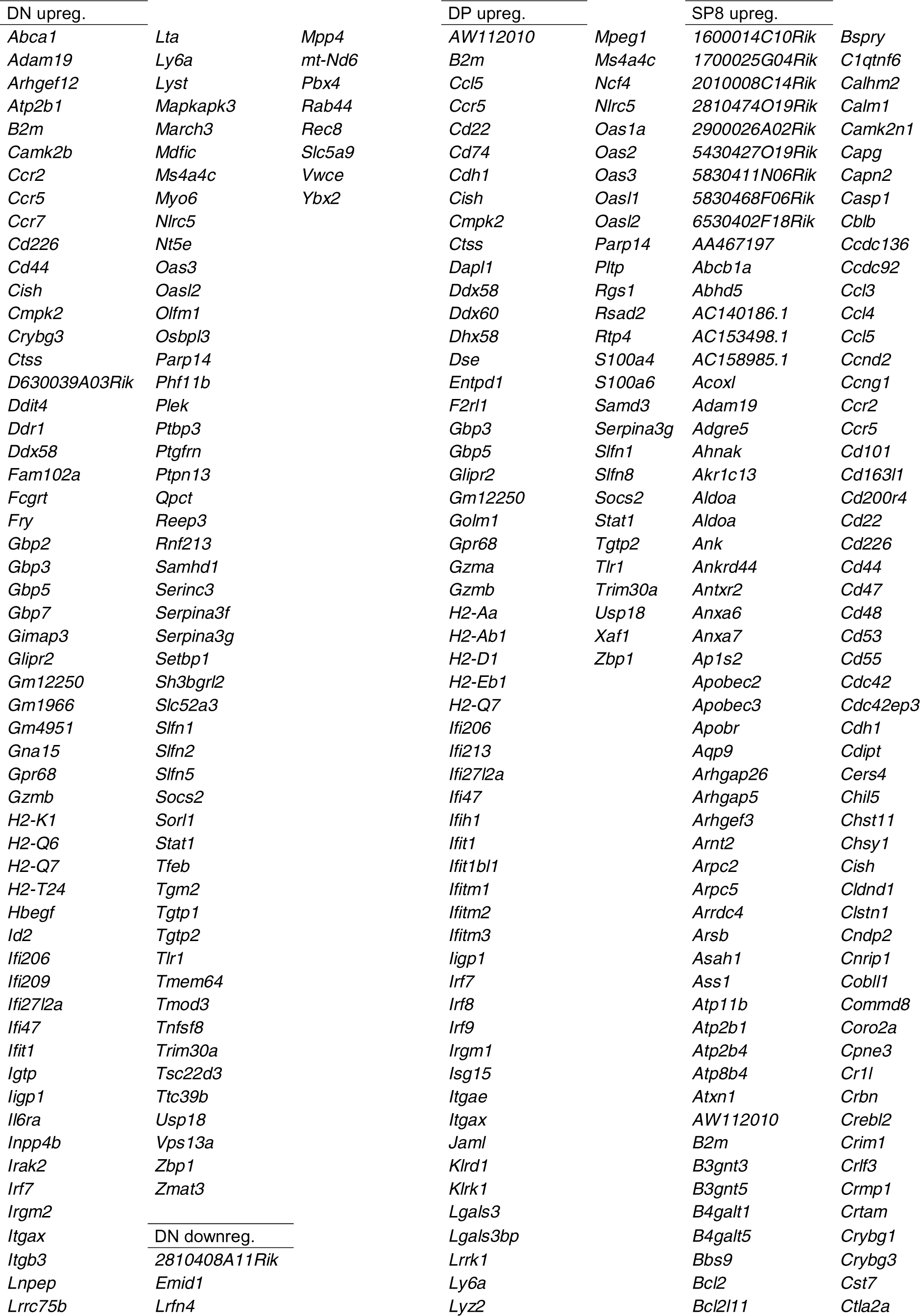
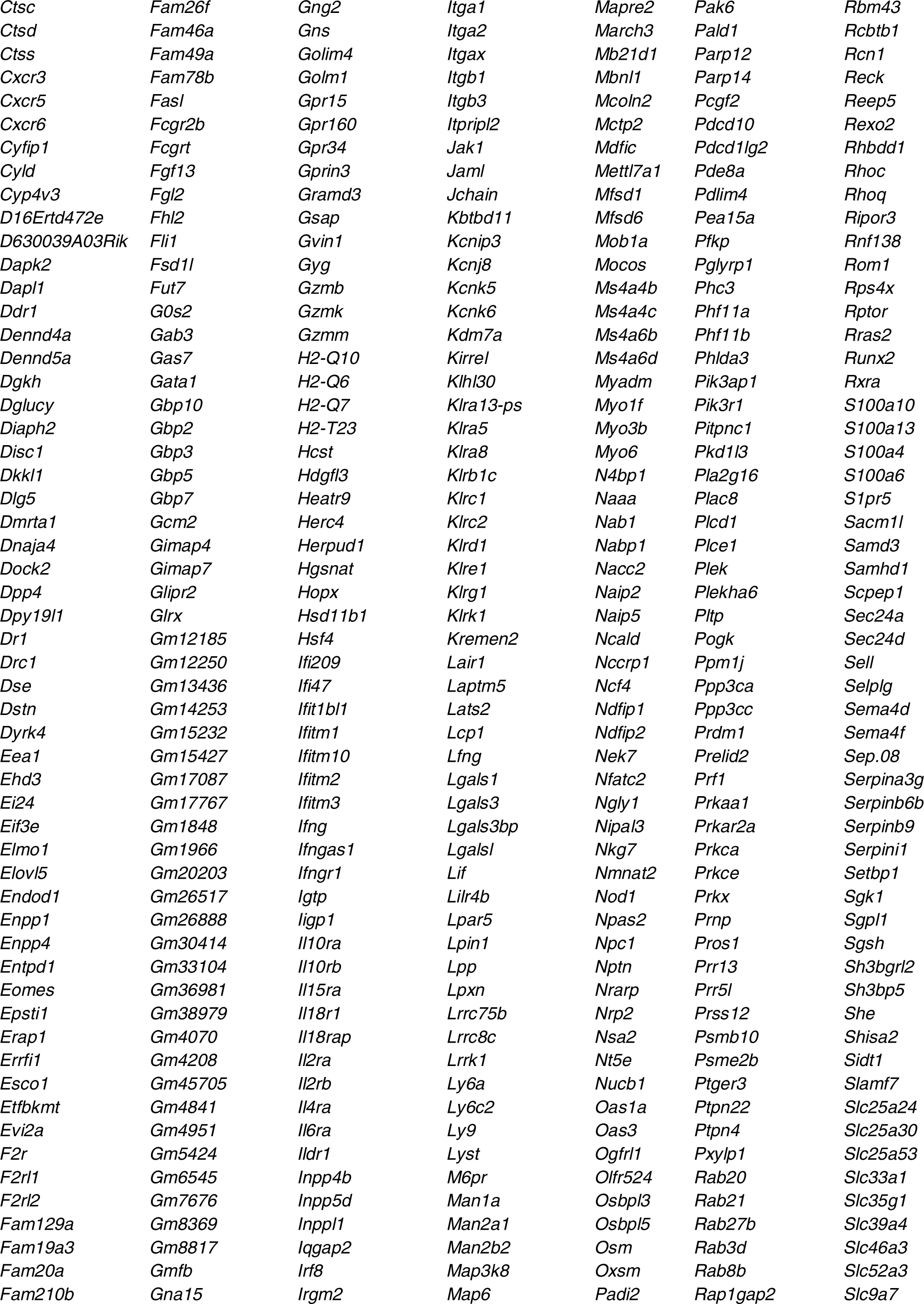
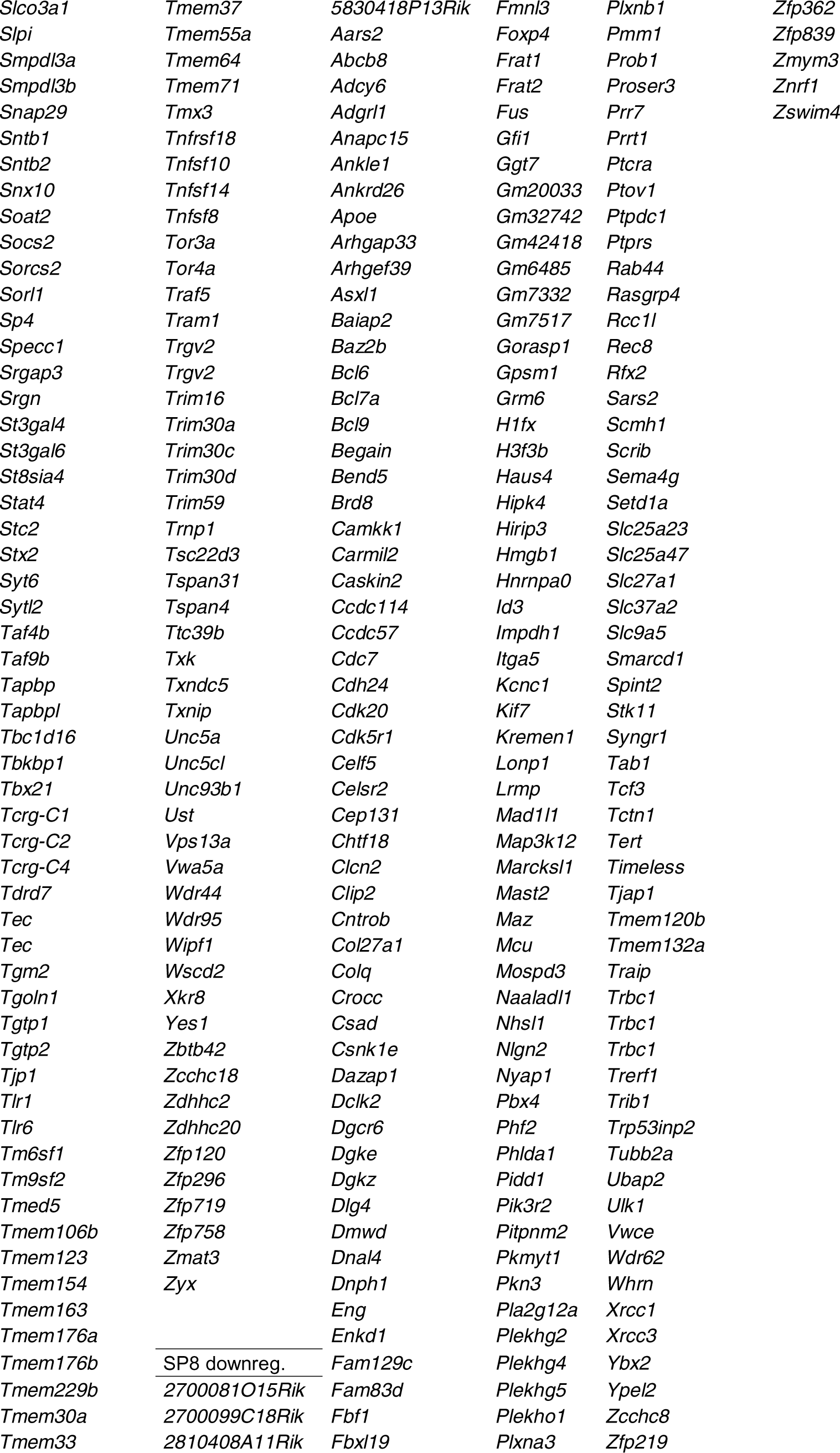
Genes that were found significantly (*Padj*<0.01) up- or downregulated mutually between STAT5B^N642H^ and STAT5A^S710F^ vs. WT mice in DN, DP or SP8 cells.

**Supplementary Table 3.**
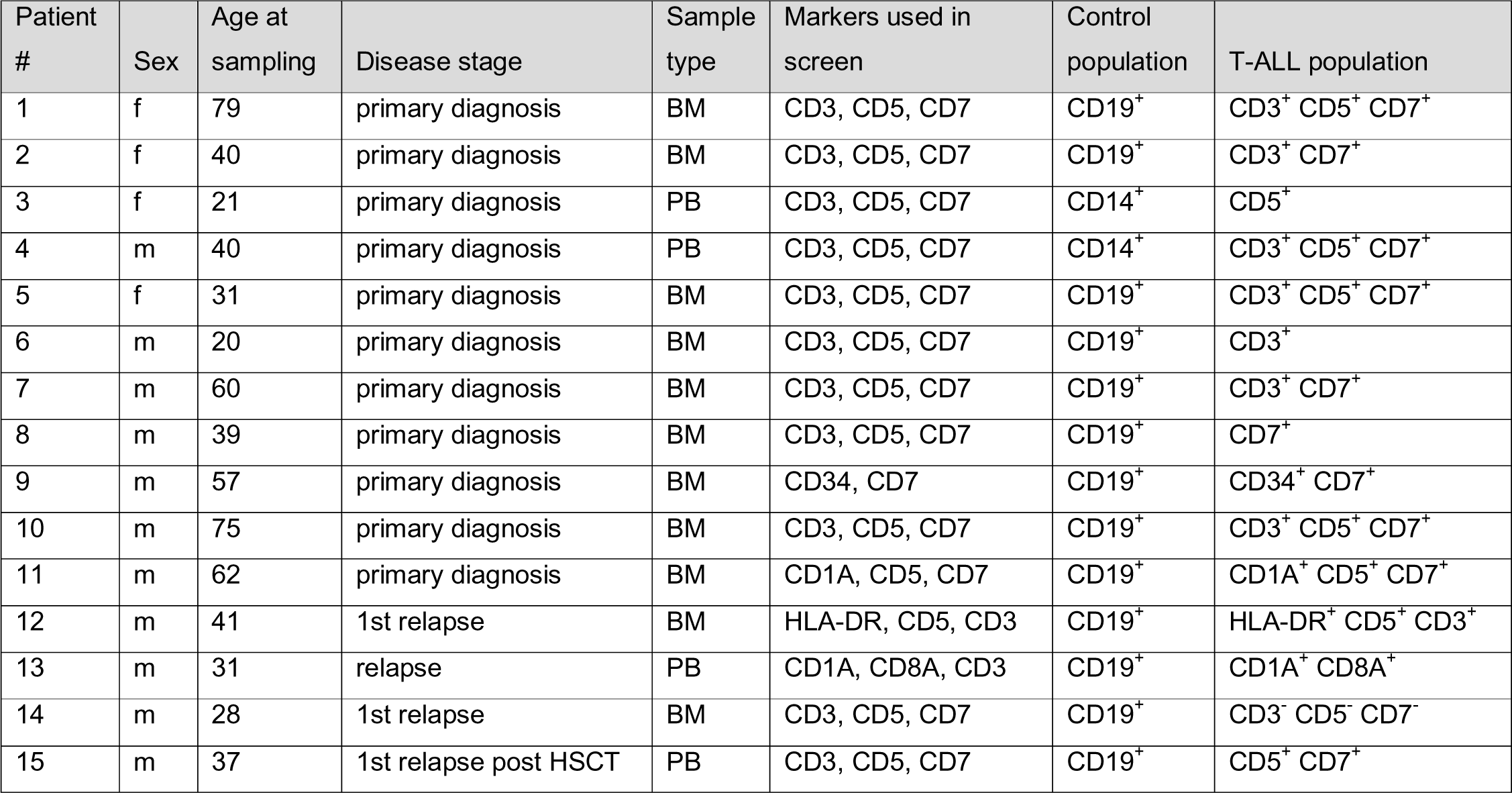
Additional clinical information on T-ALL patients, indicating sex, age and disease stage (HSCT: hematopoietic stem cell transplantation) of the patients, sample origin (BM: bone marrow, PB: peripheral blood), surface markers that were used for the screen and populations defining healthy control cells and T-ALL cells.

| Patient # | Sex | Age at sampling | Disease stage | Sample type | Markers used in screen | Control population | T-ALL population |
| --- | --- | --- | --- | --- | --- | --- | --- |
| 1 | f | 79 | primary diagnosis | BM | CD3, CD5, CD7 | CD19<sup>+</sup> | CD3<sup>+</sup> CD5<sup>+</sup> CD7<sup>+</sup> |
| 2 | f | 40 | primary diagnosis | BM | CD3, CD5, CD7 | CD19<sup>+</sup> | CD3<sup>+</sup> CD7<sup>+</sup> |
| 3 | f | 21 | primary diagnosis | PB | CD3, CD5, CD7 | CD14<sup>+</sup> | CD5<sup>+</sup> |
| 4 | m | 40 | primary diagnosis | PB | CD3, CD5, CD7 | CD14<sup>+</sup> | CD3<sup>+</sup> CD5<sup>+</sup> CD7<sup>+</sup> |
| 5 | f | 31 | primary diagnosis | BM | CD3, CD5, CD7 | CD19<sup>+</sup> | CD3<sup>+</sup> CD5<sup>+</sup> CD7<sup>+</sup> |
| 6 | m | 20 | primary diagnosis | BM | CD3, CD5, CD7 | CD19<sup>+</sup> | CD3<sup>+</sup> |
| 7 | m | 60 | primary diagnosis | BM | CD3, CD5, CD7 | CD19<sup>+</sup> | CD3<sup>+</sup> CD7<sup>+</sup> |
| 8 | m | 39 | primary diagnosis | BM | CD3, CD5, CD7 | CD19<sup>+</sup> | CD7<sup>+</sup> |
| 9 | m | 57 | primary diagnosis | BM | CD34, CD7 | CD19<sup>+</sup> | CD34<sup>+</sup> CD7<sup>+</sup> |
| 10 | m | 75 | primary diagnosis | BM | CD3, CD5, CD7 | CD19<sup>+</sup> | CD3<sup>+</sup> CD5<sup>+</sup> CD7<sup>+</sup> |
| 11 | m | 62 | primary diagnosis | BM | CD1A, CD5, CD7 | CD19<sup>+</sup> | CD1A<sup>+</sup> CD5<sup>+</sup> CD7<sup>+</sup> |
| 12 | m | 41 | 1st relapse | BM | HLA-DR, CD5, CD3 | CD19<sup>+</sup> | HLA-DR<sup>+</sup> CD5<sup>+</sup> CD3<sup>+</sup> |
| 13 | m | 31 | relapse | PB | CD1A, CD8A, CD3 | CD19<sup>+</sup> | CD1A<sup>+</sup> CD8A<sup>+</sup> |
| 14 | m | 28 | 1st relapse | BM | CD3, CD5, CD7 | CD19<sup>+</sup> | CD3<sup>-</sup> CD5<sup>-</sup> CD7<sup>-</sup> |
| 15 | m | 37 | 1st relapse post HSCT | PB | CD3, CD5, CD7 | CD19<sup>+</sup> | CD5<sup>+</sup> CD7<sup>+</sup> |

## Figure Legends

**Supplementary Figure S1.**
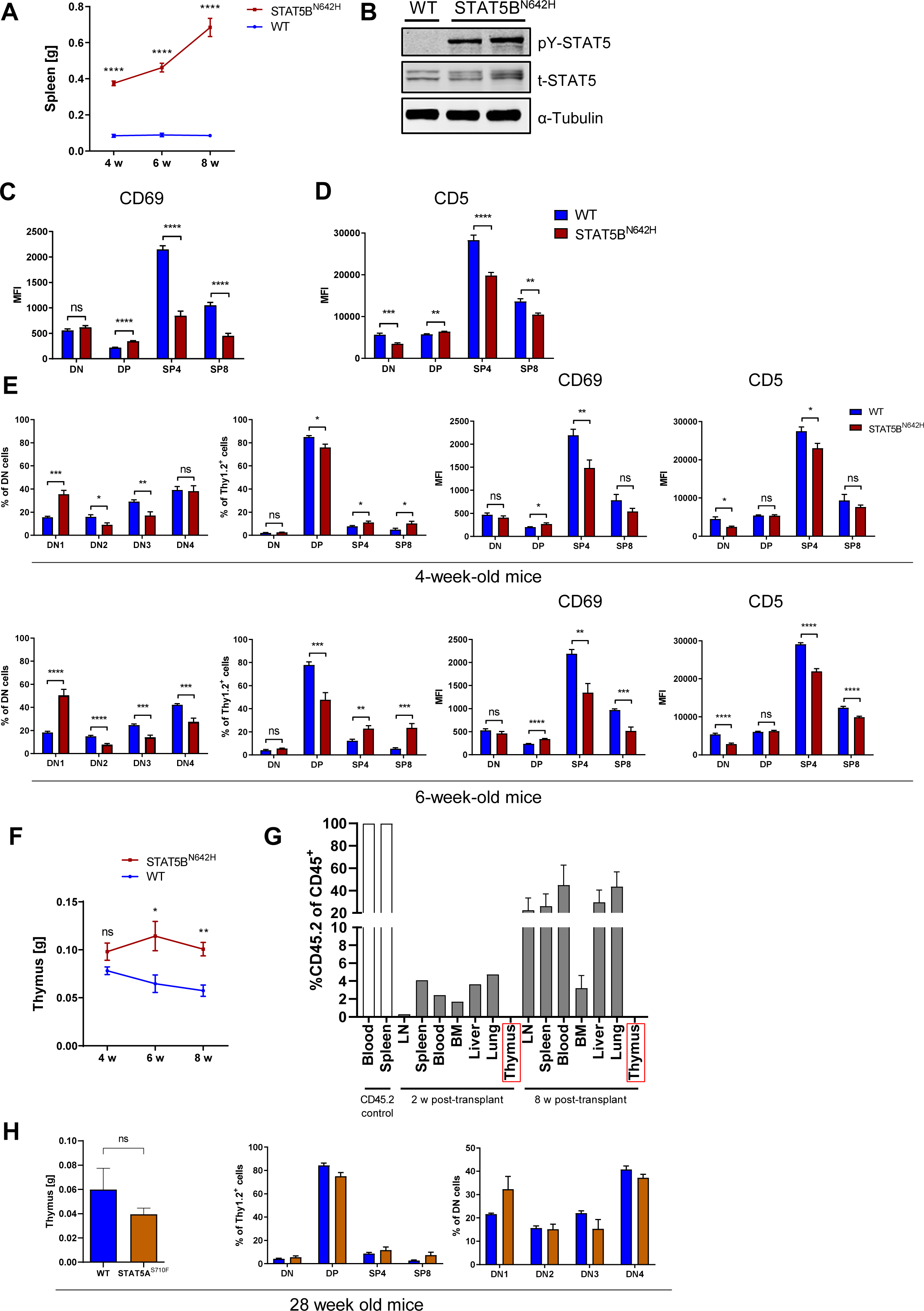
STAT5B^N642H^ but not STAT5A^S710F^ causes alterations in thymic development. **A**, Spleen weights in g from four-, six- and eight-week-old STAT5B^N642H^ or WT littermate mice. **B**, Western blot analysis for pY-STAT5, t-STAT5 expression and Tubulin as loading control in thymi from WT and STAT5^N642H^ mice. **C** and **D**, Mean fluorescent intensity (MFI) of CD69 and CD5 in DN, DP, SP4 and SP8 cells **E**, DN1, DN2, DN3, DN4 (CD25/CD44) an DN, DP, SP4, SP8 (CD4/CD8) cells gated on Thy1.2^+^ thymocytes of four- (upper panel) and six- (lower panel) week-old mice and MFI of CD69 and CD5 in DN, DP, SP4, SP8 stages. **F**, Thymus weights in g from four-, six- and eight-week-old STAT5B^N642H^ or WT littermate mice, n for all groups = 5 or more, experiments were performed twice independently. **G**, Percentages of CD45.2^+^ cells in hematopoietic organs of CD45.1 recipient mice, 2 (n = 1) or 8 weeks (n = 3) after intravenous transplantation of sorted CD8^+^ cells from LNs of terminally diseased STAT5B^N642H^ transgenic mice on CD45.2 background. Blood and spleen cells from a CD45.2 mouse were used as positive control. **H**, Thymus weights in g of 13- and 28-week-old STAT5A^S710F^ mice or WT littermates and flow cytometry analysis and percentages of DN1, DN2, DN3, DN4 (CD25/CD44) and DN, DP, SP4, SP8 (CD4/CD8) cells from these mice. **A**, **C**, **D**, **E** and **F**, Significant differences are indicated as \**P* < 0.05, \*\**P* < 0.01, \*\*\**P* < 0.001, \*\*\*\**P* < 0.0001, by two-way Anova (**A** and **F**) or unpaired two-tailed Student’s t-test (**C**, **D** and **E**). Error bars show mean +/-SEM.

**Supplementary Figure S2.**
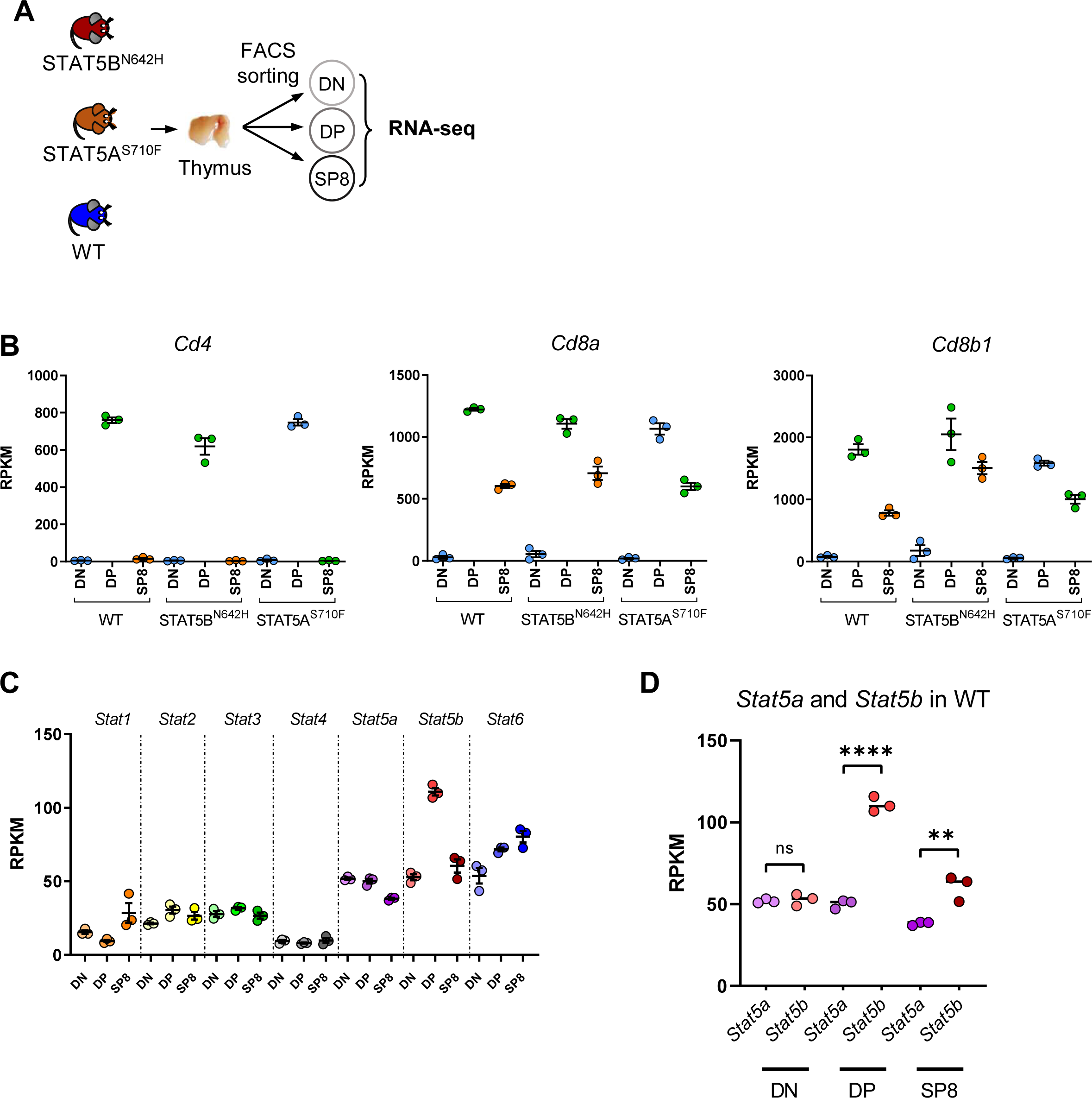
Validation of RNA-seq sample purity and GO-term analysis. **A**, Schematic depiction of thymocyte sample acquisition and RNA-seq in biological triplicates. Single cell suspensions of thymi were sorted for DN, DP and SP8 T-cell progenitor subsets by flow cytometry and subsequently subjected to RNA-seq. **B**, RPKM values of *Cd4*, *Cd8a* and *Cd8b1* confirming purity of sorted DN, DP and SP8 cells from WT, STAT5B^N642H^ and STAT5A^S710F^ thymi. **C**, RPKM values of all STAT family members in WT DN, DP and SP8 thymocytes. **D**, *Stat5a* and *Stat5b* mRNA expression in WT thymocytes. In **C**, Significant differences are indicated as **P < 0.01, ****P < 0.0001, by unpaired two-tailed Student’s t-test.

**Supplementary Figure S3.**
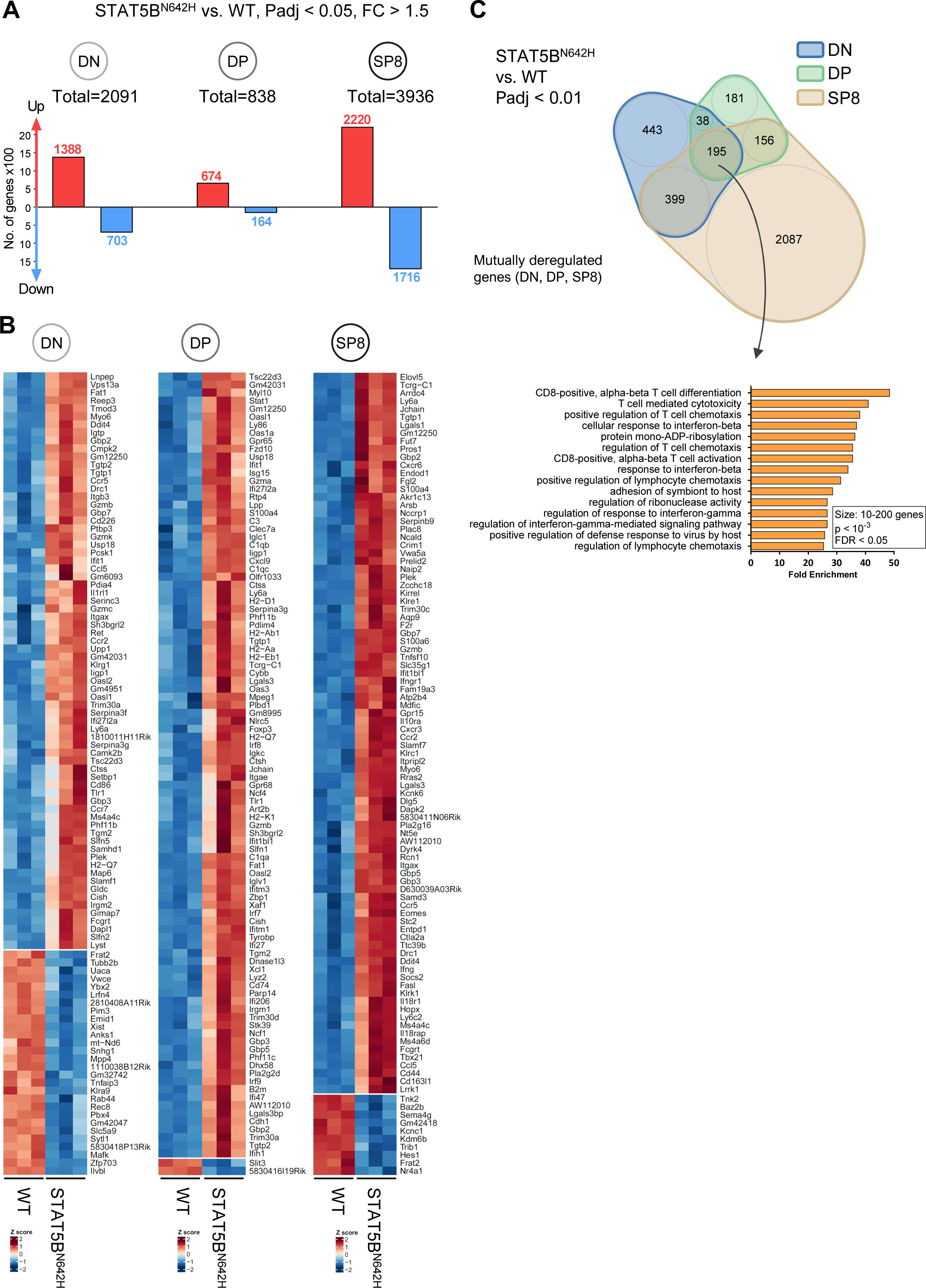
STAT5B^N642H^ leads to to high transcriptional activity and an activated T-cell phenotype. **A**, Number of up- or downregulated genes in DN, DP and SP8 cells of STAT5B^N642H^ vs. WT thymocytes (*Padj*<0.05, fc>1.5). **B**, Heatmap of the top 100 deregulated genes of these comparisons, log10 transformed normalized counts from DESeq2 were used for this analysis. **C**, Top: Venn diagram showing deregulated genes (STAT5B^N642H^ vs. WT) in DN, DP and SP8 T-cells and overlaps between subgroups, only genes with *Padj*<0.01 from respective DESeq comparisons were taken into analysis. Bottom: fold enrichment values from GO-term analysis of 195 mutually deregulated genes between all subsets, GO-terms containing 10-200 genes, *P* <10^-3^ and false discovery rate (FDR) <0.05 as inclusion criteria were taken into analysis.

**Supplementary Figure S4.**
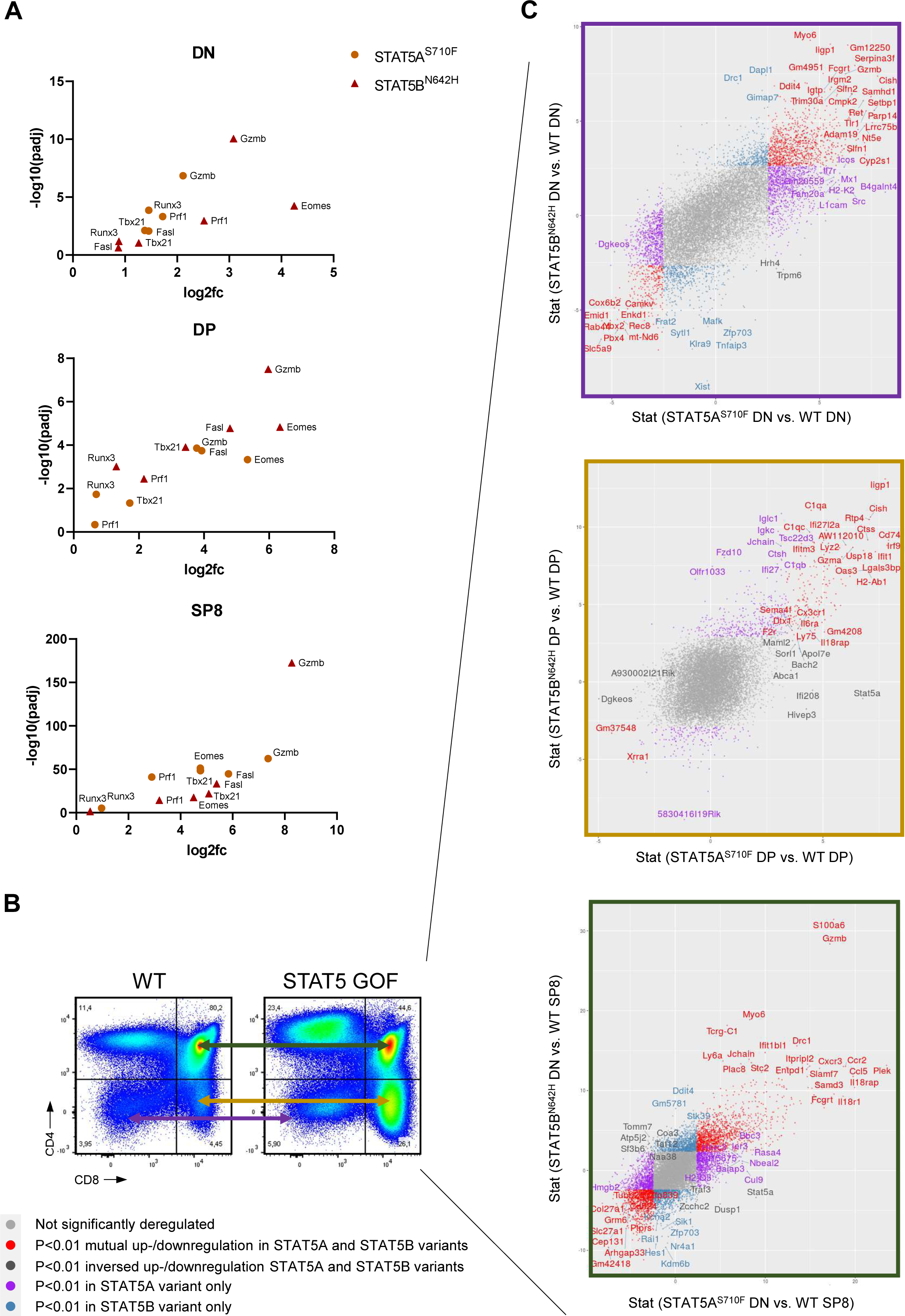
STAT5B^N642H^ and STAT5A^S710F^ mice share similar transcriptional deregulation in the thymus. **A**, *Padj* and log2fc of genes involved in T-cell activation or lineage commitment in STAT5B^N642H^ and STAT5A^S710F^ thymic subsets. Values determined by DESeq analysis. **B**, Representative flow cytometry plots indicating comparisons of thymic subsets by RNA-seq. **C**, Scatterplots showing Stat-values determined by DESeq analysis of deregulated genes of STAT5B^N642H^ vs. WT on the y-axes and STAT5A^S710F^ vs. WT on the x-axes in respective thymic subsets, as indicated in **B**.

**Supplementary Figure S5.**
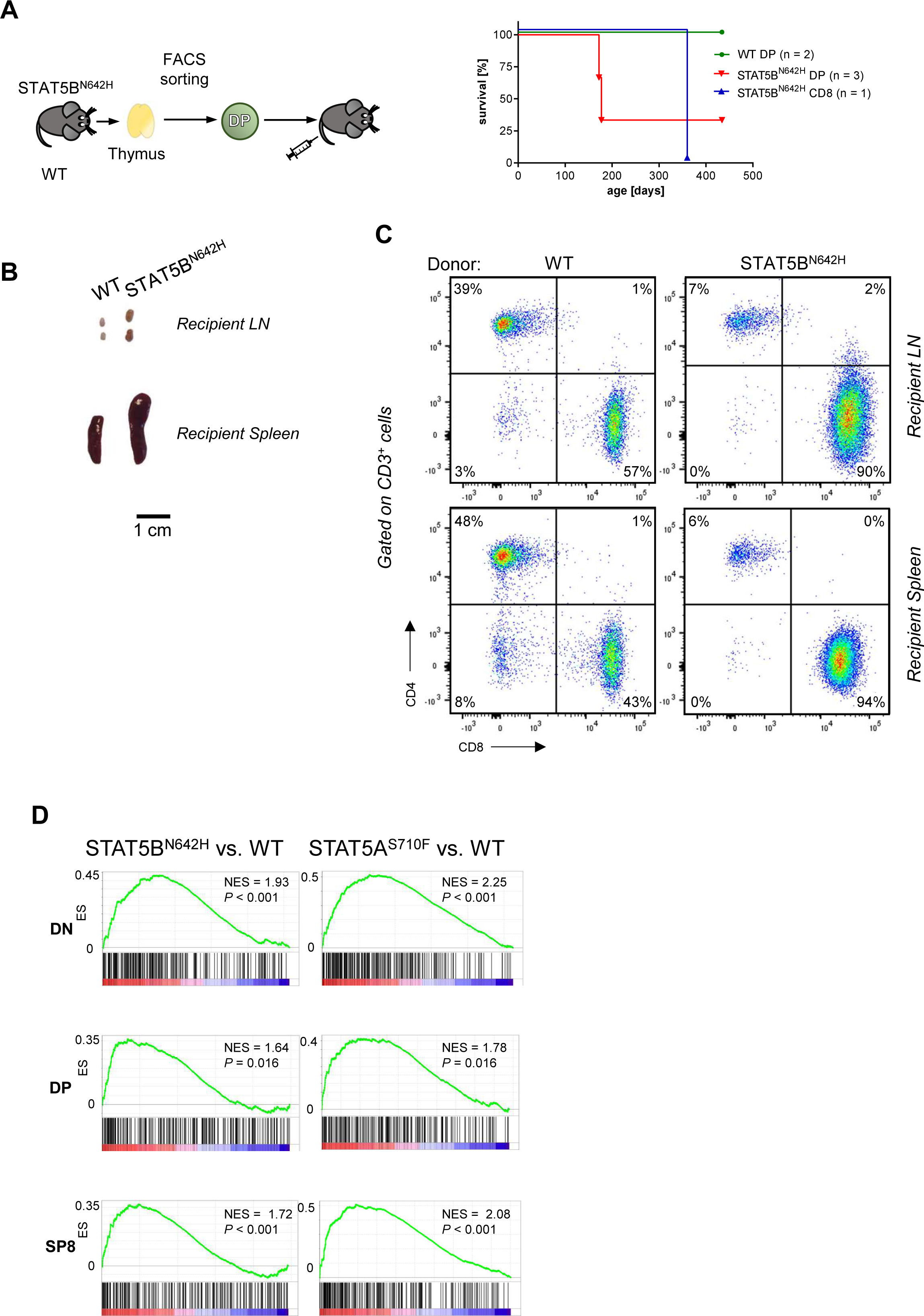
Transplanted DP cells from thymi of STAT5B^N642H^ mice give rise to CD8^+^ mature T-cell lymphoma. **A**, Schematic representation and Kaplan-Meier event-free survival plot of recipient mice transplanted with DP thymocytes from WT or STAT5B^N642H^ donor mice or CD8^+^ T-cells from LNs of STAT5B^N642H^ mice as positive control. **B**, Representative images of spleens and LNs of DP thymocyte recipients at terminal stage. **C**, CD4 and CD8 surface expression by flow cytometry analysis, gated on CD3^+^ cells of LNs and spleens from recipients of DP thymocytes. **D**, GSEA comparing top 250 upregulated genes in human ETP-ALL (10) to deregulated genes in STAT5B^N642H^ RAG2^-/-^ DN, DP or SP8 vs. respective WT populations, NES: normalized enrichment score.

**Supplementary Figure S6.**
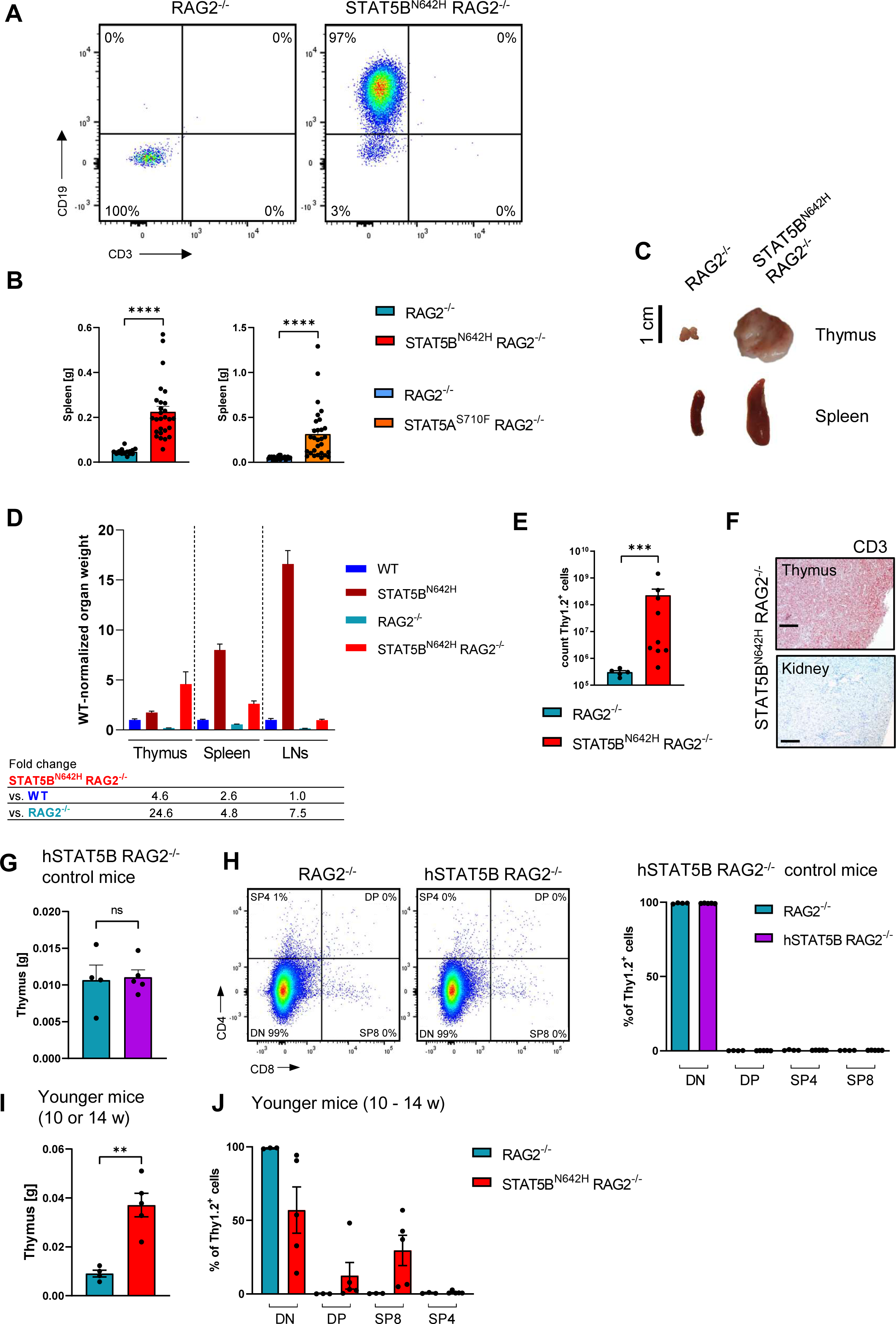
STAT5 GOF mutations induce immature T-cell neoplasia in RAG2^-/-^ background. **A**, Flow cytometry plots showing CD19 and CD3 surface expression in the LN of one STAT5B^N642H^ RAG2^-/-^ mouse that developed a B-cell lymphoma and a RAG2^-/-^ mouse as control. **B**, Spleen weights in g of diseased STAT5B^N642H^ RAG2^-/-^ (n = 27) or STAT5A^S710F^ RAG2^-/-^ (n = 29) vs. respective RAG2^-/-^ littermates (n = 17/24). **C**, Representative images of thymi and spleens of RAG2^-/-^ and diseased STAT5B^N642H^ RAG2^-/-^ mice. **D**, Relative organ weights (thymus, spleen, LN) of WT, STAT5B^N642H^, RAG2^-/-^ and STAT5B^N642H^ RAG2^-/-^ normalized to WT. Fold changes of organ weights of STAT5B^N642H^ RAG2^-/-^ compared to WT or RAG2^-/-^ are shown below, n ≥ 7 for all organs/genotypes **E**, Absolute cell counts staining positive for T-cell marker Thy1.2 in thymi of STAT5B^N642H^ RAG2^-/-^ (n = 9) and RAG2^-/-^ littermates (n = 5). **F**, Representative immunohistochemical CD3 staining in the thymus of STAT5B^N642H^ RAG2^-/-^ mice and kidney as negative control tissue, original magnification: 10x, scale bar = 200 µm. **G**, Thymus weights in g of WT human (h)STAT5B RAG2^-/-^ mice and RAG2^-/-^ littermates. **H**, Representative flow cytometry plots and analysis of CD4/CD8 staining gated on Thy1.2^+^ cells in RAG2^-/-^ and WT human (h)STAT5B RAG2^-/-^ mice. **I**, Thymus weights in g of non-terminal-diseased STAT5^N642H^ RAG2^-/-^ mice and RAG2^-/-^ littermates of 10-14 weeks of age.**J**, Representative flow cytometry plots and analysis of CD4/CD8 staining gated on Thy1.2^+^ cells in non-terminal-diseased STAT5^N642H^ RAG2^-/-^ mice and RAG2^-/-^ littermates of 10-14 weeks of age. In **B**, **E** and **I**, significant differences are indicated as \*\**P* < 0.01, \*\*\**P* < 0.001, \*\*\*\**P* < 0.0001, by a Mann-Whitney U test. Error bars show mean +/-SEM.

**Supplementary Figure S7.**
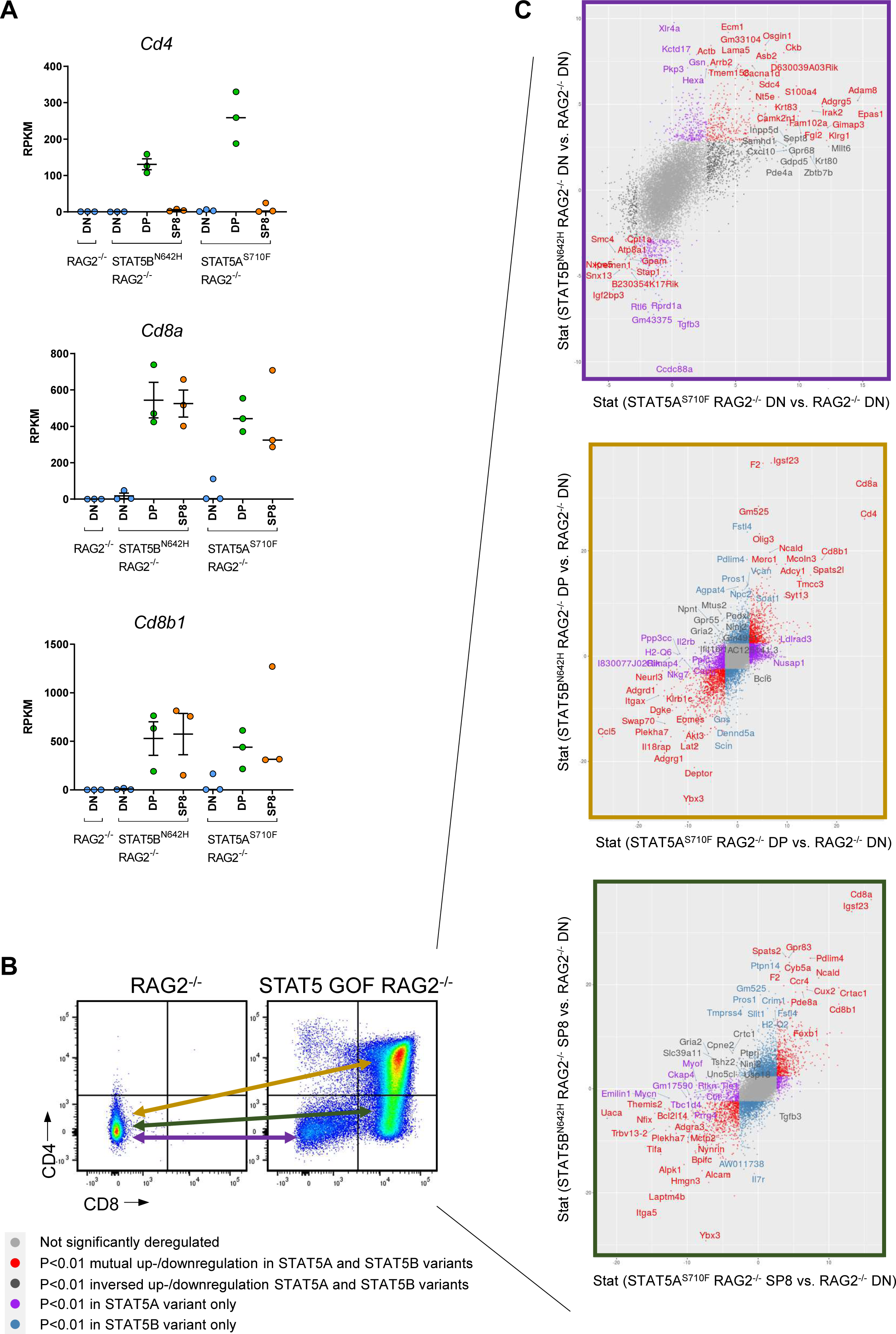
Thymic neoplasms of STAT5B^N642H^ RAG2^-/-^ and STAT5A^S710F^ RAG2^-/-^ mice display similar transcriptomic changes. **A**, RPKM values of *Cd4*, *Cd8a* and *Cd8b1* confirming purity of sorted DN, DP and SP8 cells from RAG2^-/-^, STAT5B^N642H^ RAG2^-/-^ and STAT5A^S710F^ RAG2^-/-^ thymi. **B**, Representative flow cytometry plots indicating comparisons of thymic subsets by RNA-seq. **C**, Scatterplots showing Stat-values determined by DESeq of deregulated genes of STAT5B^N642H^ RAG2^-/-^ vs RAG2^-/-^ on the y-axes and STAT5A^S710F^ RAG2^-/-^ vs. RAG2^-/-^ on the x-axes in respective thymic subsets, as indicated in **B**.

**Supplementary Figure S8.**
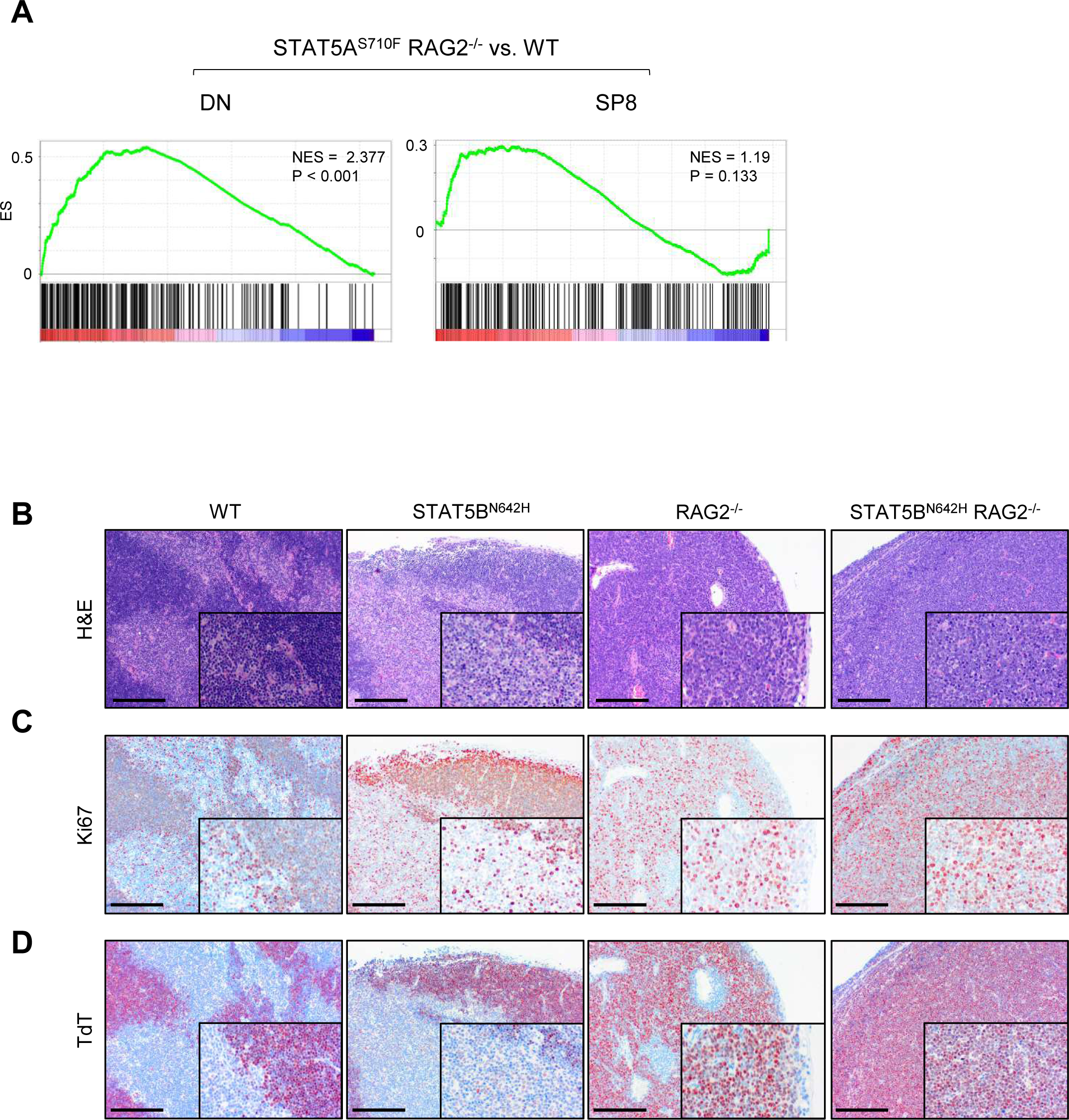
STAT5B^N642H^ RAG2^-/-^ thymic neoplasm phenotypically resemble human T-ALL. **A**, GSEA comparing top 250 upregulated genes in human ETP-ALL (10) to deregulated genes in STAT5A^S710F^ RAG2^-/-^ DN or SP8 vs. respective WT populations, NES: normalized enrichment score. Due to low transcriptional association with ETP-ALL genes, DP cells were not displayed in the analysis. **B**-**D**, Histological analysis of thymi from WT, STAT5B^N642H^, RAG2^-/-^ and STAT5B^N642H^ RAG2^-/-^ mice, staining for H&E (**B**), anti-Ki67 (**C**) and TdT (**D**), representatives of at least 3 biological replicates, original magnifications: 10x and 20x (insets), scale bar = 500 µm.

**Supplementary Figure S9.**
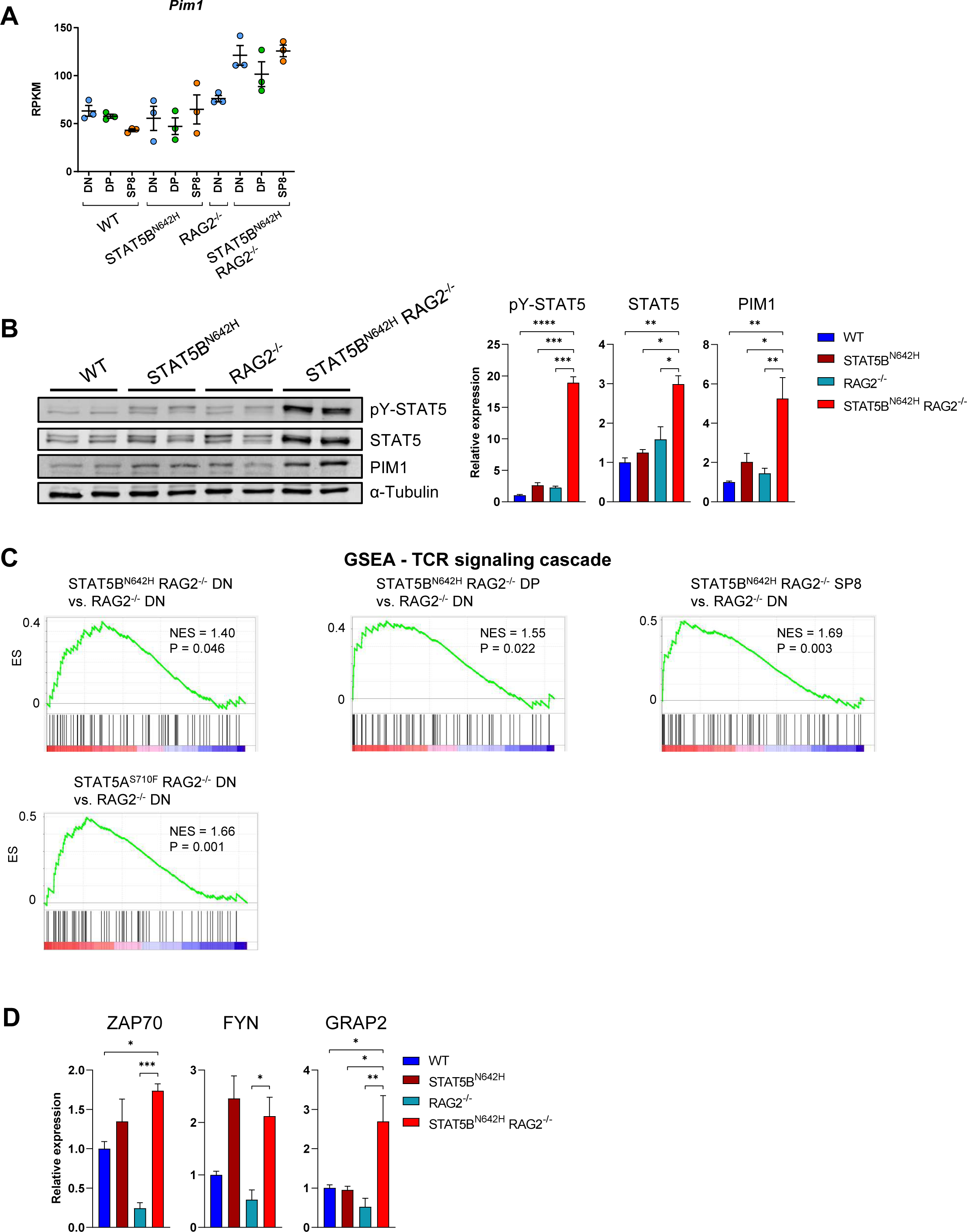
STAT5B^N642H^ RAG2^-/-^-induced neoplasms express highly activated STAT5 and its downstream oncogene PIM1. **A**, RPKM values of *Pim1* from RNA-seq in DN, DP and SP8 cells of WT, STAT5B^N642H^, RAG2^-/-^ and STAT5B^N642H^ RAG2^-/-^ thymi. **B**, Western blot analysis for pY-STAT5, t-STAT5 and PIM1 expression from whole cell extracts of thymocytes from WT, STAT5B^N642H^, RAG2^-/-^ and STAT5B^N642H^ RAG2^-/-^ mice (n = 2 each genotype) and quantification thereof. **C**, GSEA comparing indicated contrasts to 70 genes downstream of pre-TCR signaling, NES: normalized enrichment score. **D**, Relative signals to loading controls obtained from Western blot analysis for ZAP70, FYN and GRAP2 expression in thymi of mice of indicated genotypes (n = 4 each genotype). **B** and **D**, Significant differences are indicated as \**P* < 0.05, \*\**P* < 0.01, \*\*\**P* < 0.001, \*\*\*\**P* < 0.0001 by one-way Anova with Dunnet’s multiple comparison test. Error bars show mean +/-SEM.

**Supplementary Figure S10.**
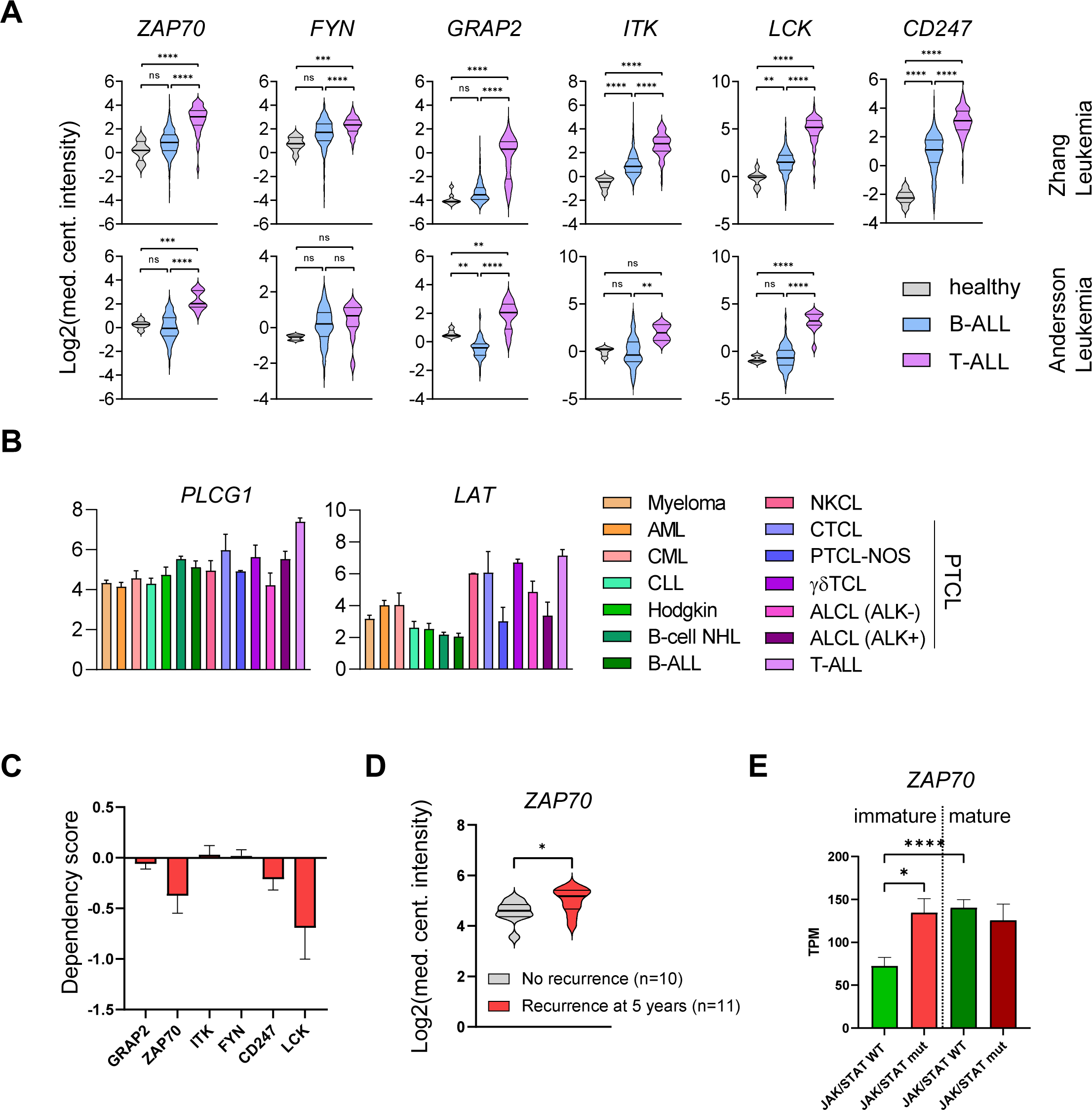
TCR pathway genes are highly expressed in human T-ALL. **A**, *ZAP70*, *FYN*, *GRAP2*, *ITK*, *CD247* and *LCK* mRNA expression data of human patients suffering from B-ALL or T-ALL and healthy bone marrow control cells. Data extracted from the Andersson Leukemia and Zhang Leukemia studies of the Oncomine database. **B**, Sequencing data showing *PLCG1* and *LAT* mRNA expression in human hematopoietic cancer cell lines. TPM: transcripts per million. **C**, Dependency scores determined by CRISPR-knockouts of indicated genes in T-ALL cell lines. Data for **B** and **C** extracted from DepMap (www.depmap.org). **D**, *ZAP70* expression of T-ALL patients with no relapse vs. patients with relapse after 5 years. Data extracted from the Oncomine database. **E**, *ZAP70* expression in TPM in immature or mature T-ALL patients, with or without activating mutations in *IL7R*, *JAK1*, *JAK3*, *STAT5A* or *STAT5B*. In **D**, significant differences are indicated as *P < 0.05, by a Mann-Whitney U test. In **A** and **E**, significant differences are indicated as *P < 0.05, \*\**P* < 0.01, \*\*\**P* < 0.001, ****P < 0.0001 by one-way Anova with Dunnet’s multiple comparison test. Error bars show mean +/-SEM.

**Supplementary Figure S11.**
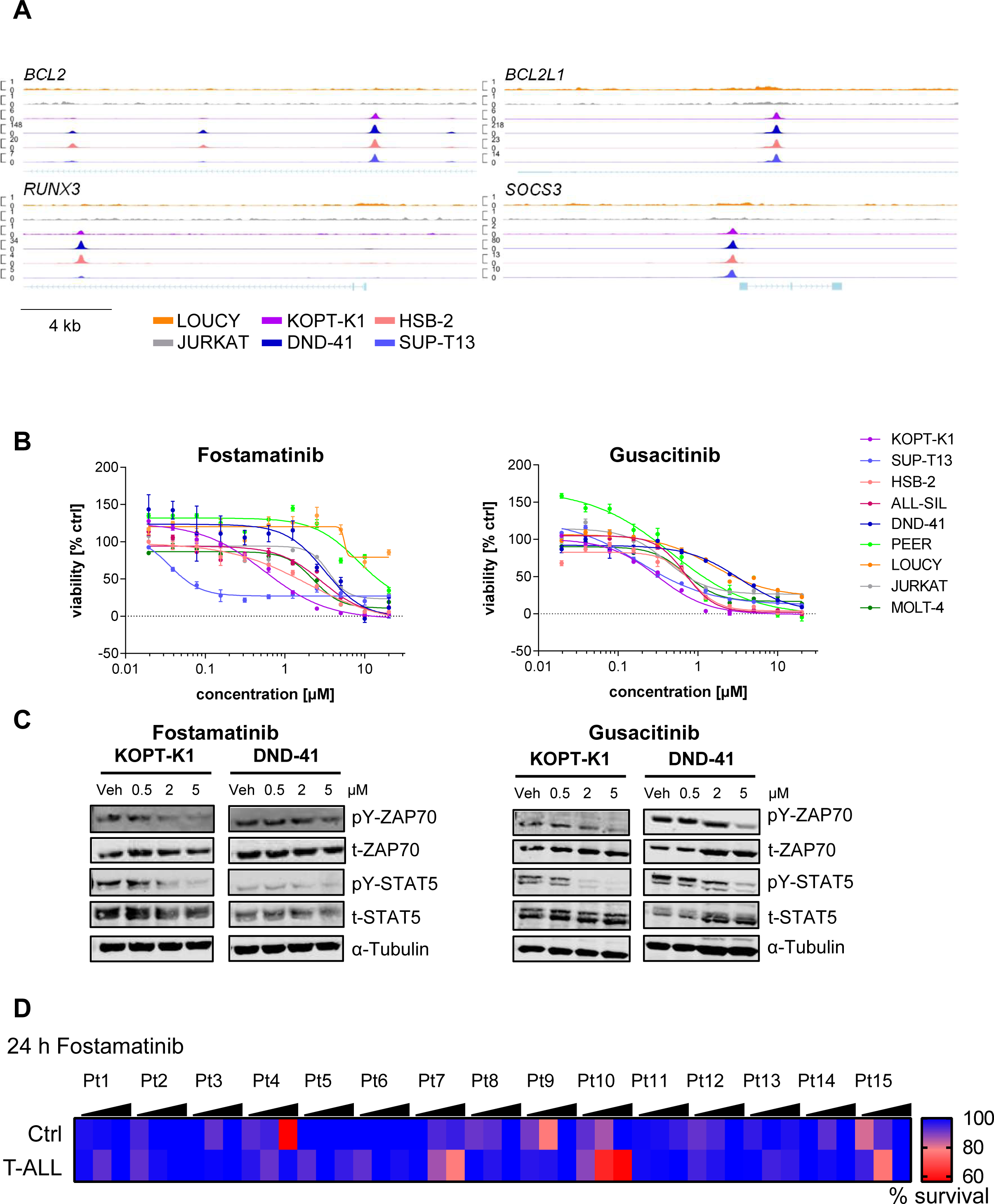
STAT5 binds to *bona fide* target genes in cell lines with high STAT5 activation and its target genes represent valuable targets in T-ALL cells. **A,** ChIP-seq lanes indicating STAT5B binding at the promoter region of genes that were previously confirmed as STAT5 target genes as indicated in 6 T-ALL cell lines, CPM indicated on the Y-axis. **B**, Representative dose-response curve for T-ALL cell lines treated with Gusacitinib or Fostamatinib at indicated concentrations determined using Cell-TiterBlue viability assay. Three independent experiments in technical triplicates were performed. Error bars show mean +/-SEM. **C**, Western blot analysis of KOPT-K1 and DND-41 cells, treated with DMSO or 0.5, 2 or 5 µM Gusacitinib or Fostamatinib for 24 h, showing expression of pY-ZAP70, t-ZAP70, pY-STAT5 and STAT5, α-Tubulin served as loading control. One representative out of two independent experiments is shown. **D**, Percent survival of healthy control and T-ALL cells of primary patient samples treated 24 h with Fostamatinib at 100, 1000 or 10,000 nM. Mean values of respective concentration vs. DMSO control in 2 technical replicates are shown. Relapsed patients are marked in red.

