## Supplementary Methods for "STAT5 Gain-of-Function Variants Promote Precursor T-Cell Receptor Activation to Drive T-Cell Acute Lymphoblastic Leukemia"

**RNA-seq data processing and analysis**

Library preparation and sequencing were performed at the Center for Molecular Medicine (CeMM), Vienna, Austria. Libraries were prepared using the NEBNext ®Ultra^TM^RNA Library Prep Kit. Next, the samples were multiplexed and sequenced on an Ilumina Hisat2000 sequencer. Quality control was performed using FastQC (version 0.11.8). Removal of low-quality bases and adapter trimming was achieved with trimmomatic, version 0.35 [1], followed by mapping using STAR aligner, version 2.7 [2], with the GRCm38 genome ([ftp.ebi.ac.uk/pub/databases/gencode/Gencode_mouse/release_M25/GRCm38.primary_assembly.genome.fa.gz](https://ftp.ebi.ac.uk/pub/databases/gencode/Gencode_mouse/release_M25/GRCm38.primary_assembly.genome.fa.gz)) and ensembl gene annotation release 93 ([ftp.ensembl.org/pub/release-93/gtf/mus_musculus/Mus_musculus.GRCm38.93.gtf.gz](http://ftp.ensembl.org/pub/release-93/gtf/mus_musculus/Mus_musculus.GRCm38.93.gtf.gz)). The genome for mapping with STAR was generated with the command:

STAR --runMode genomeGenerate --genomeDir STAR93 –genomeFastaFiles GRCm38.primary_assembly.genome.fa –sjdbGTFfile Mus_musculus.GRCm38.93.gtf.

For further analysis, only uniquely mapped reads were considered by using bamtools [3] with the command:

bamtools filter -tag NH:1 <mapped.bam> <filtered.bam>.

For RPKM calculations, the length of a gene was determined by summing up the width of the exons of a gene using the methods sum() and width(). Differential expression analysis contrasting each pair of conditions was performed using DESeq2 [4]. Gene ontology (GO) term enrichment was assessed for “biological process” using the GO Enrichment Analysis Tool at <http://geneontology.org> [5,6]. For GSEA gene lists obtained from DESeq2 analysis were ranked by fold change and subjected to the GSEAPreranked tool provided by the Broad Institute, version GSEA_Linux_4.0.2. The gene sets from single-cell RNA-seq data of human T-cells were taken from Szabo et al., 2019 [7]. The ETP-ALL gene sets were obtained as follows: microarray data (Table_S24_12SJETP_40SJnonETP_limma.xlsx) from Zhang et al. 2012 [8] was filtered for probes with refseq ID. The top upregulated genes between ETP-ALL and non-ETP-ALL were determined by ranking via the T-statistic column. Top ranking microarray probes that target different regions of the same gene were not double counted. Accordingly, the number of top probes was set to yield the desired number of genes. Thus, the top 352 microarray probes cover 250 genes. The TCR signaling gene set was compiled from the MSigDB and literature research.

The Venn Diagram was visualized with nvennR [9]. Differentially expressed genes with *P*<0.01 in STAT5B^N642H^ vs. WT contrasts of DN, DP and SP8 cells were intersected. Up- and downregulated genes common to all contrasts were subjected to GO analysis (see above).

**Immunohistochemistry and histologic analysis**

Mouse organs were incubated 24 h in 4% phosphate-buffered formaldehyde solution (Roti-Histofix; Carl Roth) on a rotator at 4°C, dehydrated, embedded, and cut (4-μm-thick sections). For immunohistochemical staining, heat-mediated antigen retrieval was performed in citrate buffer at pH 6.0 (Dako) and stained with antibodies against CD3 (Thermo Fisher Scientific; RM-9107-S0; dilution 1:300), Ki67 (Novocastra, Leica Biosystem; NCL-Ki67p; dilution 1:1000), TdT (eBioscience; 14-9739-82; dilution 1:50) and pY-ZAP70 (Cell Signaling Technology, 2701, 1:400) using standard protocols. Images were taken using an Olympus BX 53 LED light microscope with an Olympus SC50 camera or a Leica DMi8 with a Leica DMC 2900 camera. Images were analyzed using the ImageJ (version 1.53a) software.

**ChIP-seq data processing and analysis**

Preprocessing of the raw sequence reads was done with PRINSEQ-lite [10] (version 0.20.4). The remaining high quality reads were aligned with BWA [11] (version 0.7.15-r1140) against the human reference genome (GRCh38) and further processed to bam files with samtools [12] (version 1.4). The deepTools [13] (version 3.5.1) bamCompare function was applied to create CPM normalized bigWig files. These were then visualized in R [14] (version 4.2.1) with rtracklayer [15] (version 1.56.1). ChIP-seq peaks were called with MACS27 (version 2.1.0) against the respective input controls.

| **FACS antibodies (for murine cells)** | |  |  |  |  |  |  |
| --- | --- | --- | --- | --- | --- | --- | --- |
| **Target** | **Fluorochrome** | **Clone** | | | | **Company** | **Cat. no.** |
| CD19 | eFluor450 | eBio1D3 | | | | Invitrogen | 48-0193-82 |
| CD25 | APC | PC61.5 | | | | Invitrogen | 17-0251-82 |
| CD3E | FITC | 145-2C11 | | | | Biolegend | 100306 |
|  | PerCP-Cyanine5.5 | 145-2C11 | | | | Invitrogen | 45-0031-82 |
| CD4 | APC-eFluor 780 | RM4-5 | | | | Invitrogen | 47-0042-82 |
|  | FITC | GR1.5 | | | | Invitrogen | 11-0041-82 |
|  | PE-Cyanine7 | GK1.5 | | | | Invitrogen | 25-0041-82 |
| CD44 | PE | IM7 | | | | Invitrogen | 12-0441-82 |
| CD5 | PE-Cyanine7 | 53-7.3 | | | | Invitrogen | 25-0051-81 |
| CD69 | FITC | H1.2F3 | | | | Invitrogen | 11-0691-82 |
| CD8A | PerCP-Cyanine5.5 | 53-6.7 | | | | Invitrogen | 45-0081-82 |
|  | PE | 53-6.7 | | | | Invitrogen | 12-0081-82 |
| Ly5.1 | PE | A20 | | | | Invitrogen | 12-0453-82 |
| Ly5.2 | APC | 104 | | | | Invitrogen | 17-0454-82 |
| Ter119 | PE | TER-119 | | | | Invitrogen | 12-5921-81 |
| Thy1.2 | eFluor450 | 53-2.1 | | | | Invitrogen | 48-0902-82 |
|  | APC | 53-2.1 | | | | Invitrogen | 17-0902-81 |
| Viability dye | APC-eFluor780 |  | | | | Invitrogen | 65-0865-14 |
| **FACS antibodies (for human cells)** | |  | | | |  |  |
| **Target** | **Fluorochrome** | **Clone** | | | | **Company** | **Cat. no.** |
| CD1a | PE-Cy7 | HI149 | | | | Biolegend | 300122 |
|  | APC | HI149 | | | | BD Biosciences | 559775 |
| CD3 | PE | HIT3a | | | | Biolegend | 300308 |
| CD5 | APC-Cyanine7 | L17F12 | | | | Biolegend | 364010 |
| CD7 | FITC | 6B7 | | | | Biolegend | 343104 |
|  | PE-Cyanine7 | 6B7 | | | | Biolegend | 343114 |
| CD8A | APC | SK1 | | | | Biolegend | 344722 |
| CD14 | APC | PC61.5 | | | | Biolegend | 301808 |
| CD19 | APC | HIB19 | | | | Biolegend | 302211 |
|  | FITC | SJ25C1 | | | | Biolegend | 363008 |
| CD34 | APC | 8G12 | | | | BD Biosciences | 1196918 |
| Viability dye | DAPI |  | | | | Biolegend | 422801 |

**Supplementary Methods Table 1.** Antibodies and dyes used for FACS measurement and sorting, Cat. no.: catalogue number

| **Western blot antibodies** | |  |  |  |
| --- | --- | --- | --- | --- |
| **Target** | **Dilution** | **Clone** | **Company** | **Cat. no.** |
| STAT5 | 1:1000 | 89/Stat5 | BD | 610191 |
| BCL-2 | 1:1000 | 124 | Cell Signaling Technology | 15071S |
| PIM1 | 1:1000 | D8D7Y | Cell Signaling Technology | 54523S |
| ZAP70 | 1:1000 | 521626 | R&D Systems | MAB3709 |
| pY-STAT5 (Tyr694/699) | 1:1000 | Polyclonal | Invitrogen | 71-6900 |
| pY-ZAP70 (Tyr319) | 1:1000 | 65E4 | Cell Signaling Technology | 2717 |
| pY-SRC (Tyr416) | 1:1000 | D49G4 | Cell Signaling Technology | 6943 |
| pY-PLCγ1 (Tyr783) | 1:1000 | D6M9S | Cell Signaling Technology | 14008 |
| α-Tubulin | 1:5000 | DM1A | Santa Cruz Biotechnology | sc-32293 |
| Actin | 1:5000 | Polyclonal | Santa Cruz Biotechnology | C-11 |

**Supplementary Methods Table 2.** Antibodies used for Western blot analysis, Cat. no.: catalogue number

| **ChIP antibodies** | |  |
| --- | --- | --- |
| **Target** | **Company** | **Cat. no.** |
| STAT5B | R&D Systems | AF1584 |
| STAT5B | Invitrogen | 13-5300 |

**Supplementary Methods Table 3.** Antibodies used for ChIP-seq analysis, Cat. no.: catalogue number
